# Functional analysis of *Salix purpurea* genes support roles for *ARR17* and *GATA15* as master regulators of sex determination

**DOI:** 10.1101/2023.04.21.537837

**Authors:** Brennan Hyden, Dana L. Carper, Paul E. Abraham, Guoliang Yuan, Tao Yao, Leo Baumgart, Yu Zhang, Cindy Chen, Ronan O’Malley, Jin-Gui Chen, Xiaohan Yang, Robert L. Hettich, Gerald A. Tuskan, Lawrence B. Smart

## Abstract

The Salicaceae family is of growing interest in the study of dioecy in plants because the sex determination region (SDR) has been shown to be highly dynamic, with differing locations and heterogametic systems between species. Without the ability to transform and regenerate *Salix* in tissue culture, previous studies investigating the mechanisms regulating sex in the genus *Salix* have been limited to genome resequencing and differential gene expression, which are mostly descriptive in nature, and functional validation of candidate sex determination genes has not yet been conducted. Here we used Arabidopsis to functionally characterize a suite of previously identified candidate genes involved in sex determination and sex dimorphism in the bioenergy shrub willow *Salix purpurea*. Six candidate master regulator genes for sex determination were heterologously expressed in Arabidopsis, followed by floral proteome analysis. In addition, 11 transcription factors with predicted roles in mediating sex dimorphism downstream of the SDR were tested using DAP-Seq in both male and female *S. purpurea* DNA. The results of this study provide further evidence to support models for the roles of *ARR17* and *GATA15* as master regulator genes of sex determination in *S. purpurea*, contributing to a regulatory system that is notably different from that of its sister genus *Populus*. Evidence was also obtained for the roles of two transcription factors, an *AP2*/*ERF* family gene and a homeodomain-like transcription factor, in downstream regulation of sex dimorphism.

## Introduction

Understanding the genetic regulation of sex determination in dioecious plants is of interest in the plant biology community because, while dioecy is only observed in about six percent of angiosperm species, it is present in numerous taxa and families and is thought to have evolved independently on as many as 5,000 occasions (Käfer et al., 2017). As such, understanding the genetic mechanisms that lead to separation of sexes in different taxa can provide insight into the repeated evolution and maintenance of dioecy. The Salicaceae family is of particular interest in this effort, as nearly all species in the family are dioecious, and it contains two genera of economic importance: poplars (*Populus*) and willows (*Salix*). In particular, *Salix* contains over 300 species, which are native to every continent except Antarctica and Australia, and grow in a diverse range of biomes, including subarctic tundra, deserts and temperate and tropical forests (Argus, 1997; Kuzovkina et al., 2007). *Salix* species also exhibit a variety of growth habits, ranging from prostrate dwarfs to shrubs and trees (Argus, 1997). Despite this remarkable diversity in species range and form, dioecy has been maintained throughout the evolution of most of the family, including all *Populus* and *Salix*. Moreover, the sex determination region (SDR) in *Salix* appears to have shifted among chromosomes and varied heterogametic systems on multiple occasions, with tree willows such as *S. nigra* and *S. chaenomeloides* containing SDR on Chr07 with an XY system (Sanderson et al., 2021; Wang et al., 2022) and alternatively on Chr15 with an XY system in *S. arbutifolia* (Wang et al., 2022). Shrub willows, including *S. purpurea* and *S. viminalis*, contain the SDR on Chr15 under a ZW system (Pucholt et al., 2015; Zhou et al., 2018; Wilkerson et al., 2022). Because of the dynamic nature of sex determination in Salicaceae, there is an opportunity to characterize the precise mechanisms of sex determination in diverse species across this family and to add to our understanding of the evolution, conservation, and transition of the SDR.

Much work has already been done to identify candidate master regulator genes in both *Populus* and *Salix*. Müller et al (2020) demonstrated that a homolog of *ARR17*, an Arabidopsis type-C response regulator, is likely the sole master regulator gene in *Populus*, with females expressing *ARR17* while in males it is either absent or silenced by smRNA produced from exon 1 repeats located on the Y chromosome (Müller et al., 2020). A similar *ARR17* mediated sex determination system is thought to be present in some willows with Chr07 and Chr15 XY systems, but has not been confirmed through expression or functional analysis (Wang et al., 2022). *ARR17* was first proposed as a candidate master regulator gene of sex determination in *S. purpurea* (Chr15 ZW SDR) by Zhou et al. (2020), where the authors identified four inverted repeats of the gene on Chr15W in a female *S. purpurea* (Zhou et al., 2020). Another study conducted RNA-Seq and smRNA-Seq of an F_2_ family of *S. purpurea* and proposed several candidate master regulator genes in addition to *ARR17*, but notably did not find evidence for the smRNA silencing mechanism in males that exists in XY *Populus* (Hyden et al., 2021). This lack of smRNA expression, along with confirmed expression of *ARR17* in both male and female *S. purpurea*, suggests that if *ARR17* is a master regulator gene in *S. purpurea*, it may operate via a mechanism that differs from *Populus* and the tree willows (Hyden et al., 2021). Based on sequencing and expression analysis, several other genes have been proposed as candidate genes for sex determination in addition to *ARR17*, including homologs of *GATA15*, *AGO4*, *DRB1*, and several hypothetical proteins (Hyden et al., 2021).

However, no functional validation has been conducted to confirm the role of these proposed sex-determination genes. Recently, sequencing and expression data from a monoecious *S. purpurea* revealed a structural variant on Chr15W that includes deletions of *ARR17*, *AGO4*, and *DRB1*, but not *GATA15* (Hyden et al., 2023). Based on these data, we hypothesize that *GATA15* is a master regulator gene of sex determination that functions to promote female development, while *ARR17* acts as a suppressor of male development.

Previous research on willow sex determination and dimorphism has relied primarily on the comparison of DNA and RNA sequencing data between males and females (Carlson et al., 2017; Zhou et al., 2020; Hyden et al., 2021); however, functional validation is ultimately needed to support these hypotheses. Yet, unlike poplar, there is not a facile protocol for *Salix* transformation and regeneration from tissue culture, so it is not feasible to study gene function in willow using transformation-based gain/loss-of function approaches. Our best alternative is to use a model plant species, such as Arabidopsis, for transgenic manipulation. In addition, transcription factors can also be studied *in vitro* using DAP-Seq as it does not require plant transformation. In DAP-Seq, transcription factors are transiently expressed and incubated with native genomic DNA. Transcription factor bound DNA fragments are then sequenced and aligned to the reference genome (O’Malley et al., 2016). This system enables transcription factor binding analysis in any species from which candidate genes can be cloned and high-quality DNA can be sequenced.

In this study, we sought to elucidate the mechanisms of sex determination and dimorphism in *S. purpurea* by investigating both master regulator genes of sex and transcription factors with predicted roles in sex dimorphism. Six candidate master regulator genes for sex determination in *S. purpurea* were heterologously expressed in Arabidopsis using a constitutive promoter. Bottom-up proteomics was performed on floral tissue of the transgenic Arabidopsis plants to identify proteins regulated by the overexpressed candidate master regulators of primary sex dimorphism (anther or stamen development). To characterize regulation of sex dimorphism downstream of the SDR, which includes floral development and floral secondary metabolite production, DAP-Seq was performed on 11 transcription factors. Nine of the TFs tested are located on autosomes and have eQTL in the *S. purpurea* SDR and two are located in the SDR and are candidate master regulator genes of sex, for which expression in Arabidopsis and proteomic analysis was also performed (Hyden et al., 2021). Using data from these combined methods, we constructed a conceptual model for the functional role of several genes involved in sex determination and dimorphism and provide evidence to support of *ARR17* and *GATA15* as master regulator genes of sex determination in *S. purpurea*.

## Results

### Proteomic Analysis

Transgenic Arabidopsis floral proteomics data were used to assess the broad impact of each candidate *S. purpurea* gene on the proteome as well as changes among proteins relevant to floral development and sex determination biological processes. Eight transgenic constructs were generated (six candidate master regulator gene and two methodological control genes), each with 4-6 independent insertion events (Table 1) and proteome data were obtained from three full-sibling T_2_ plants from each event in addition to five empty vector control plants, for a total of 122 plants evaluated. Across all 122 samples, a total of 17,191 Arabidopsis protein accessions were identified and the total proteins identified for each transgenic line ranged from 11,395 to 16,885 (Table 1). The relative impact of each heterologous expression transgenic line on proteome expression was compared against the empty vector control proteome data. Across the eight expression lines, the number of proteins of differential abundance (PDA) ranged from 103 to 5,970 (Supplemental Dataset S20). Enriched MapMan functional categories (Schwacke et al., 2019) were identified for all eight heterologously expressed genes to identify biological processes affected by expression of each candidate master regulator (Fig. S1–S2). As expected, the impact of each expressed gene on the floral proteome was not uniform, as shown by the variation in total peptides detected, PDAs, and MapMan enrichment categories. Plants expressing *GATA15* showed the greatest number of PDAs when compared against the empty vector control (Fig. 1, Table 1), followed by hypothetical protein Sapur.15WG074900 and *ARR17* with 5,318 and 4,305 PDAs respectively (Fig. 1, Fig. S3, Table 1). The positive control genes *LEAFY* and *FT* affected only 444 and 179 PDAs, respectively (Fig. S3, Table 1). The CCHC Zinc finger Sapur.15WG068800 produced the fewest number of PDAs, at only 103.

**Figure 1.**
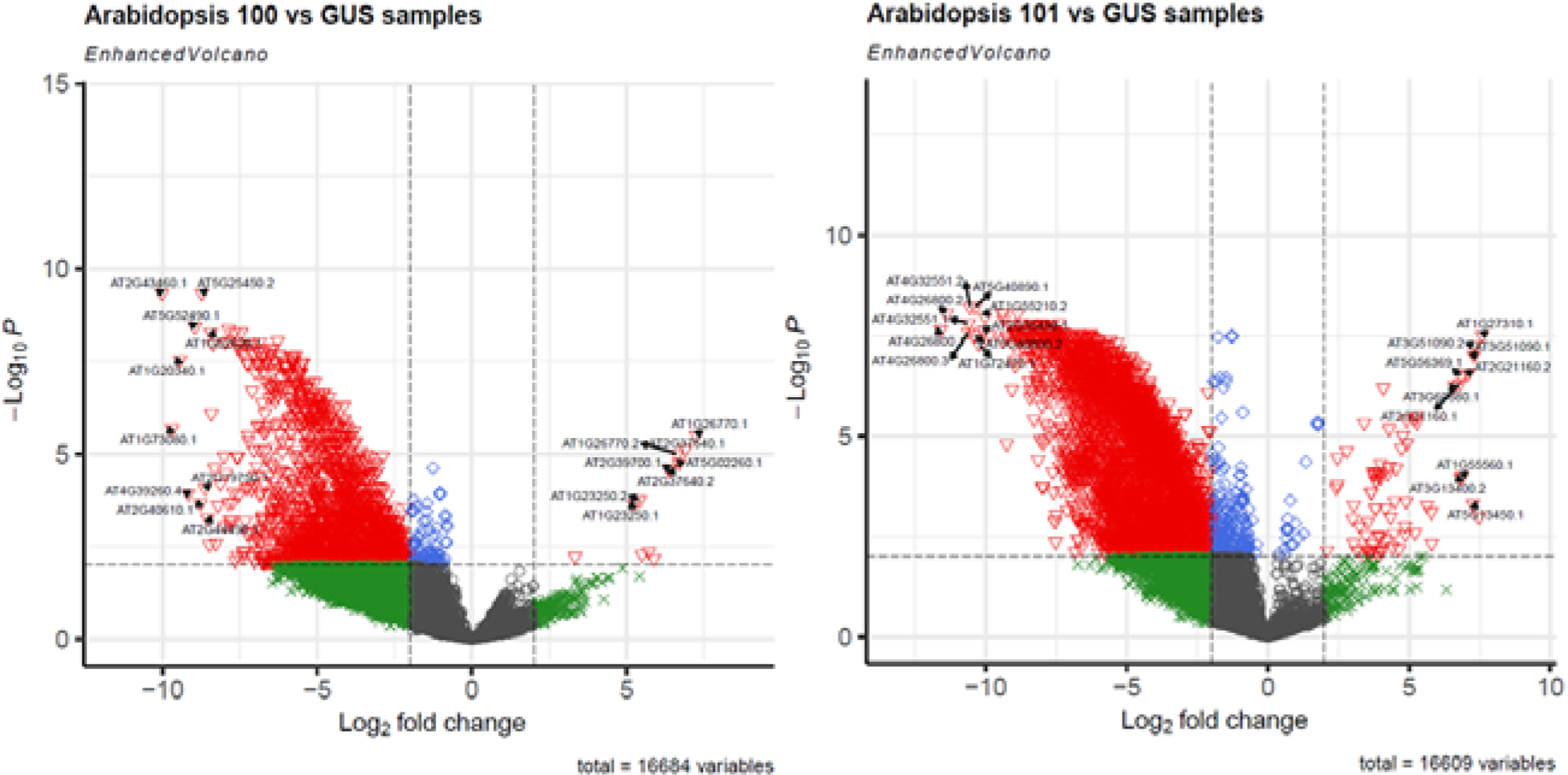
Volcano plots displaying the total proteins of differential abundance (PDA) results for the *ARR17* (100, left) and *GATA15* (101, right) expression lines, relative to the empty vector control. Grey indicates non-significant proteins, green indicates those that meet the log2 FC cutoff but not p-value, blue indicates those that meet the p-value threshold but not log2 FC, and red indicates proteins that have met the log2 FC cutoff and p-value threshold. Identities of the most extreme PDA are indicated.

**Table 1.** Total, differentially abundant proteins (PDA), and unique PDA determined for each transgenic heterologous expression line in Arabidopsis.

| Construct | Expressed Gene | Gene Annotation | Events | Total Proteins | PDA (FDR < 0.05) | Unique PDA |
| --- | --- | --- | --- | --- | --- | --- |
| pBH100 | Sapur.15WG073500 | *ARR17* response regulator | 6 | 16,684 | 4305 | 1236 |
| pBH101 | Sapur.15WG062800 | *GATA15* transcription factor | 4 | 16,609 | 5970 | 1953 |
| pBH102 | Sapur.15WG068800 | CCHC Zinc Finger | 4 | 16,023 | 103 | 45 |
| pBH103 | Sapur.15WG074300 | *DRB1* dsRNA binding | 6 | 12,306 | 343 | 110 |
| pBH104 | Sapur.15WG074900 | Hypothetical Protein | 5 | 16,885 | 5318 | 1041 |
| pBH106 | Sapur.15WG075700 | Hypothetical Protein | 4 | 11,355 | 611 | 137 |
| pBH107 | Sapur.15WG122200 | *LEAFY* | 5 | 12,271 | 444 | 65 |
| pBH108 | Sapur.008G061900 | *Flowering Locus T* | 5 | 11,928 | 179 | 57 |

In general, most transgenic proteomes contained a substantial number of downregulated proteins when compared to the empty vector control and this observation was particularly pronounced when expressing *ARR17*, *GATA15*, and Sapur.15WG074900 (Fig. 2). Resulting PDAs across each heterologous expression line were compared to assess similarity in the resulting proteome changes. Among all the PDAs, there was considerable proteome changes observed between *ARR17*, *GATA15*, and Sapur.15WG074900 hypothetical protein lines (Fig. 2). Notably, in addition to the large number of resulting PDAs, GATA15 and ARR17 expression lines exhibited unique protein abundance profiles of multiple floral development genes, indicating a likely role in floral and reproductive development and sex determination.

**Figure 2.**
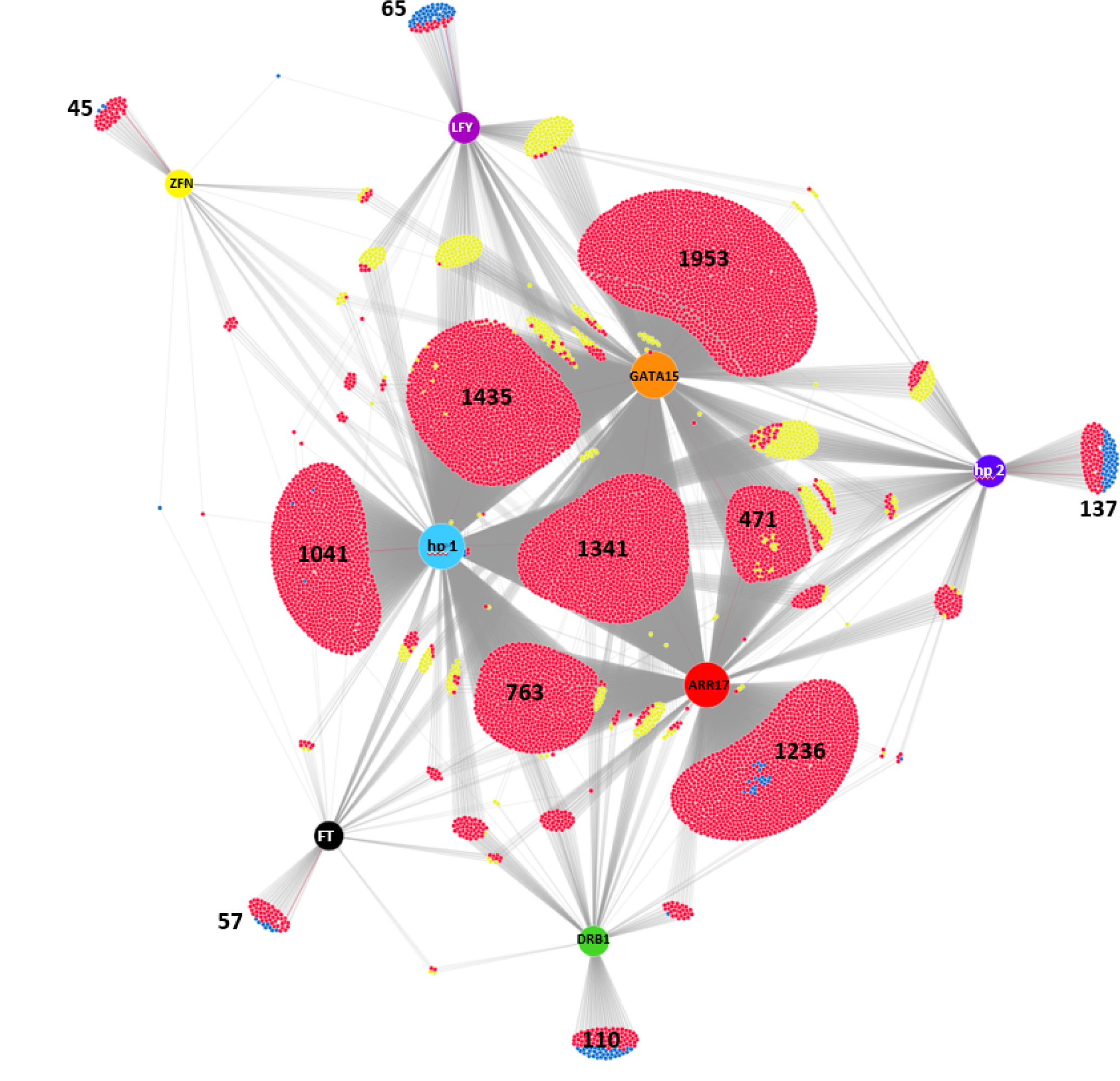
DiVenn diagram comparing proteomic expression patterns in transgenic plants heterologously expressing each of the eight *S. purpurea* genes relative to the empty vector control. Colored nodes reperesent expressed genes (ZFN: Sapur.15WG068800; hp 1: Sapur.15WG074900; hp 2: Sapur.15WG075700). Proteins are grouped according to unique differential abundance or shared differential abundance among the Arabidopsis lines. Relative to the empty vector control, red points represent downregulated PDAs, blue points are upregulated PDAs, and yellow points are PDAs which show differing abundance patterns between expression lines. Groupings containing unique PDAs and the largest groupings of shared PDA are labeled with the respective number of proteins in each group.

### Sapur.15WG072500 ARR17 expression results

The *ARR17*-OX lines (pBH100) showed 4,305 PDAs, 1,236 of which had unique abundance patterns when compared to other heterologously expressed candidate genes (Table 1, Fig. 1–2). Enriched MapMan categories included chromatin organization, coenzyme metabolism, transferase and hydrolase enzyme activity, multi-process regulation, protein homeostasis and modification, RNA biosynthesis and processing, and vesicle trafficking (Fig. S1). Among the proteins showing differential abundance unique to the *ARR17*-OX lines were several with annotations related to floral development, including a homolog of *PISTILLATA* and seven genes involved in tapetum and pollen development (Table 2). Among the proteins most regulated in the *ARR17*-OX lines were multiple expansin family proteins, which have been shown to be involved in pollen tube development and cell expansion (Table S1, Fig. 1) (Liu et al., 2021).

**Table 2.** Floral and reproductive development genes showing differential expression unique to either *ARR17* or *GATA15* expression lines. log_2_FC, log_2_ fold-change; FDR, false discovery rate

| OX Gene | Arabidopsis Gene | log2FC | FDR | Annotation |
| --- | --- | --- | --- | --- |
| ARR17 | AT1G19890.1 | 5.42 | 2.03E-02 | male-gamete-specific histone H3, *MALE-GAMETE-SPECIFIC HISTONE H3* |
|  | AT5G51860.2 | 1.80 | 4.54E-02 | *AGAMOUS*-like 72 |
|  | AT3G12145.1 | -1.02 | 4.12E-02 | *FLOR1, FLORAL TRANSITION AT THE MERISTEM* |
|  | AT2G30800.2 | -1.36 | 4.24E-02 | helicase in vascular tissue and tapetum |
|  | AT5G05560.1 | -1.87 | 4.52E-02 | pollen calcium-binding protein 1, *EMBRYO DEFECTIVE 2771*, anaphase promoting complex 1 |
|  | AT5G05560.3 | -2.01 | 2.57E-02 | pollen calcium-binding protein 1, *EMBRYO DEFECTIVE 2771*, anaphase promoting complex 1 |
|  | AT3G11980.1 | -2.75 | 2.98E-02 | *FATTY ACID REDUCTASE 2, MALE STERILITY 2* |
|  | AT5G20240.2 | -3.02 | 7.83E-03 | *PISTILLATA* |
|  | AT3G10390.3 | -3.03 | 4.59E-03 | *FLOWERING LOCUS D*, Reduced Systemic immunity 1 |
|  | AT1G67990.1 | -3.28 | 3.33E-02 | *TAPETUM-SPECIFIC METHYLTRANSFERASE 1* |
|  | AT3G10390.2 | -3.37 | 2.55E-04 | *FLOWERING LOCUS D*, Reduced Systemic immunity 1 |
|  | AT3G10390.1 | -3.54 | 1.56E-04 | *FLOWERING LOCUS D*, Reduced Systemic immunity 1 |
|  | AT3G10390.4 | -3.54 | 3.72E-04 | *FLOWERING LOCUS D*, Reduced Systemic immunity 1 |
|  | AT1G25260.1 | -6.50 | 1.43E-08 | *REDUCED POLLEN NUMBER 1, REDUCED POLLEN NUMBER* |
| GATA15 | AT3G12145.1 | 0.79 | 4.07E-04 | *FLOR1, FLORAL TRANSITION AT THE MERISTEM* |
|  | AT4G29010.1 | -1.31 | 4.93E-02 | *ABNORMAL INFLORESCENCE MERISTEM* |
|  | AT3G58780.3 | -1.37 | 1.76E-02 | *SHATTERPROOF 1, AGAMOUS*-like 1 |
|  | AT3G58780.2 | -1.42 | 2.64E-02 | *SHATTERPROOF 1, AGAMOUS*-like 1 |
|  | AT5G51860.1 | -2.43 | 4.76E-03 | *AGAMOUS*-like 72 |
|  | AT5G51860.2 | -2.45 | 1.37E-03 | *AGAMOUS*-like 72 |
|  | AT5G40260.1 | -6.42 | 1.27E-07 | *RUPTURED POLLEN GRAIN1* |
|  | AT4G32551.1 | -10.60 | 1.55E-08 | *ROTUNDA2, LEUNIG* |
|  | AT4G32551.2 | -10.55 | 6.10E-09 | *ROTUNDA2, LEUNIG* |

### Sapur.15WG062800 GATA15 expression results

The *GATA15*-OX lines (pBH101) showed the greatest number of floral PDAs by a substantial margin at 5,970, of which 1,953 had unique abundance patterns (Table 1, Fig. 1-2), as well as the greatest number of enriched MapMan functional categories, at 22. Among the enriched MapMan categories were cell cycle organization, transcriptional regulation, RNA modification, and vesicle transport (Fig. S1). *GATA15* was one of two genes tested, the other being *ARR17,* whose expression resulted in a unique differential abundance pattern of proteins with floral development annotations, including the upregulation of *FLOR1*, a gene involved in floral meristem development and transition (Acevedo et al., 2004), and downregulation of two isoforms each of *SHATTERPROOF1* homologs and *AGL72* homologs, involved in fruit dehiscence and floral transition, respectively (Liljegren et al., 2000; Dorca-Fornell et al., 2011) (Table 2). Among the proteins with the greatest abundance in the *GATA15*-OX lines were *SKU5*-Similar 13 and *SKU5*-Similar 14, the former of which has been shown to be essential for pollen tube growth through regulation of jasmonic acid biosynthesis (Zhang et al., 2022). However, these proteins do not appear to be uniquely upregulated by *GATA15*, as they were also upregulated in the hypothetical protein Sapur.15WG074900 transgenic lines. Among the most downregulated proteins in *GATA15*-OX lines were two isoforms of *LEUNIG*, which is involved in regulating gynoecium development (Table 2, Table S1, Fig. 1).

### Sapur.15WG068800 CCHC Zinc Finger Transcription Factor expression results

The *Sapur.15WG068800-OX* lines (pBH102) had the fewest PDAs relative to the control at only 103, of which 45 had unique protein abundance patterns (Table 1, Fig. S3). Enriched functions included phosphorylation, carrier-mediated transport, and solute channel transport (Fig. S2).

### Sapur.15WG074300 DRB1 expression results

The lines expressing *DRB1* (pBH103) had 343 PDAs relative to the empty vector control, of which 110 showed unique abundance patterns (Table 1, Fig. 2, Fig. S3). Enriched functional categories included sucrose metabolism, chromatin structure, hydrolase enzyme activity, and MAP kinase cascade signaling (Fig. S2).

### Sapur.15WG74900 hypothetical protein expression results

In addition to *ARR17* and *GATA15*, plants expressing the hypothetical protein Sapur.15WG074900 (pBH104) also showed an exceptionally high number of PDAs at 5,318, of which 1,041 had unique abundance patterns (Table 1, Fig. 2, Fig. S3). Despite this large number of floral PDAs, there were not any protein annotations among these that have been previously related to floral development (two *SKU5*-like proteins were upregulated, but their upregulation was also observed in the *GATA15*-OX lines). Sapur.15WG074900 lines also showed the second greatest number of enriched functional categories, with 20 that were significant (Fig. S2), including vesicle trafficking, RNA splicing, homeostasis, and modification, phosphorylation, ubiquitin-proteasome system, calcium-dependent signaling, fatty acid metabolism, and sucrose metabolism.

### Sapur.15WG075700 hypothetical protein expression results

Plants expressing the hypothetical protein Sapur.15WG075700 (pBH106) had 611 PDAs, of which 137 were unique (Table 1, Fig. S3). Microfilament network, fatty acid metabolism, protein quality control, phosphorylation, transcriptional regulation, RNA export, and primary active transport of solutes were all enriched functions in these plants (Fig. S2).

### Sapur.15WG122200 LEAFY control expression results

A *S. purpurea* homolog of *LEAFY*, located on Chr15, was expressed as a positive experimental control (pBH107). The *LEAFY*-OX lines had 444 PDAs, of which only 65 were unique (Table 1, Fig. S3). Among the significantly enriched functional terms were pectin, microtubular network, MAP kinase cascade signaling, and transcriptional regulation (Fig. S2).

### Sapur.008G061900 Flowering Locus T control expression results

A *S. purpurea Flowering Locus T* (*FT*) homolog was also included as a positive experimental control (pBH108), and showed 179 PDAs, of which 57 had a unique abundance pattern (Table 1, Fig. S3). The *FT*-OX lines showed enrichment for the fewest functional categories, with only two that were significant: oxidoreductase enzymes and transcriptional regulation (Fig. S2). Arabidopsis T_2_ progeny from all five *S. purpurea Flowering Locus T* (*FT*) expression events showed early flowering phenotypes when compared to the empty vector control, producing inflorescences in just 23 days after germination (Fig. S4). These observations are consistent with the Arabidopsis *FT* overexpression phenotype (Kardailsky et al., 1999) and indicated the *S. purpurea FT* homolog was both expressed and functional in Arabidopsis, which is the first such example of functional expression of an *S. purpurea* gene in Arabidopsis. Among the most upregulated proteins in the FT-OX lines were three isoforms of *FASCIATA5*, which is involved in floral initiation and is consistent with the role of *FT* in inducing early flowering (Fig. 1, Table S1) (Albert et al., 2015).

### Genome-wide identification of transcription factor binding sites

In DAP-Seq, peak calling is performed by mapping transcription factor bound DNA fragments to the reference genome and comparing the relative read abundance to background levels of mapping, with a three-fold mapping rate relative to background being a standard minimum cutoff for identifying putative transcription factor binding sites, also termed “summits”. Candidate target genes are identified as those nearest to a summit with expression in the direction away from the summit (i.e. antisense if upstream of the summit, positive sense if downstream of the summit). Of the 11 transcription factors tested by DAP-Seq, three (Sapur.001G003600.1 (*AP2/ERF*, GCCGGC binding sequence), Sapur.003G027300.1 (Homeodomain-like, TGGATAA binding sequence), and Sapur.15WG062800.1 (*GATA15*, GATCA binding sequence)) produced large numbers of significant peaks with a threshold of three-fold or greater mapping in the 94006 female and ‘Fish Creek’ male libraries along with similar binding motif predictions in both libraries (Table 3, Fig. S5). In particular, Sapur.001G003600.1 had 12 peaks in 94006 and five peaks in ‘Fish Creek’ with a mapping rate at least 10-fold over background, while Sapur.003G027300.1 had 36 in 94006 and 56 in ‘Fish Creek’. The remaining eight transcription factors that were tested produced inconsistent motif predictions between the two libraries and fewer than 20 significant peaks in any library (Table 3, Fig. S5, Supplemental Dataset S21). Many of the peaks were shared among these eight transcription factors, suggesting that the results from these latter eight genes are likely the result of background mapping and not true transcription factor binding sites. All three genes that produced prominent binding motifs and more than 20 significant binding sites also targeted multiple floral development genes (Table S2), confirming a likely role in the regulation of primary sex dimorphism and development.

**Table 3.**
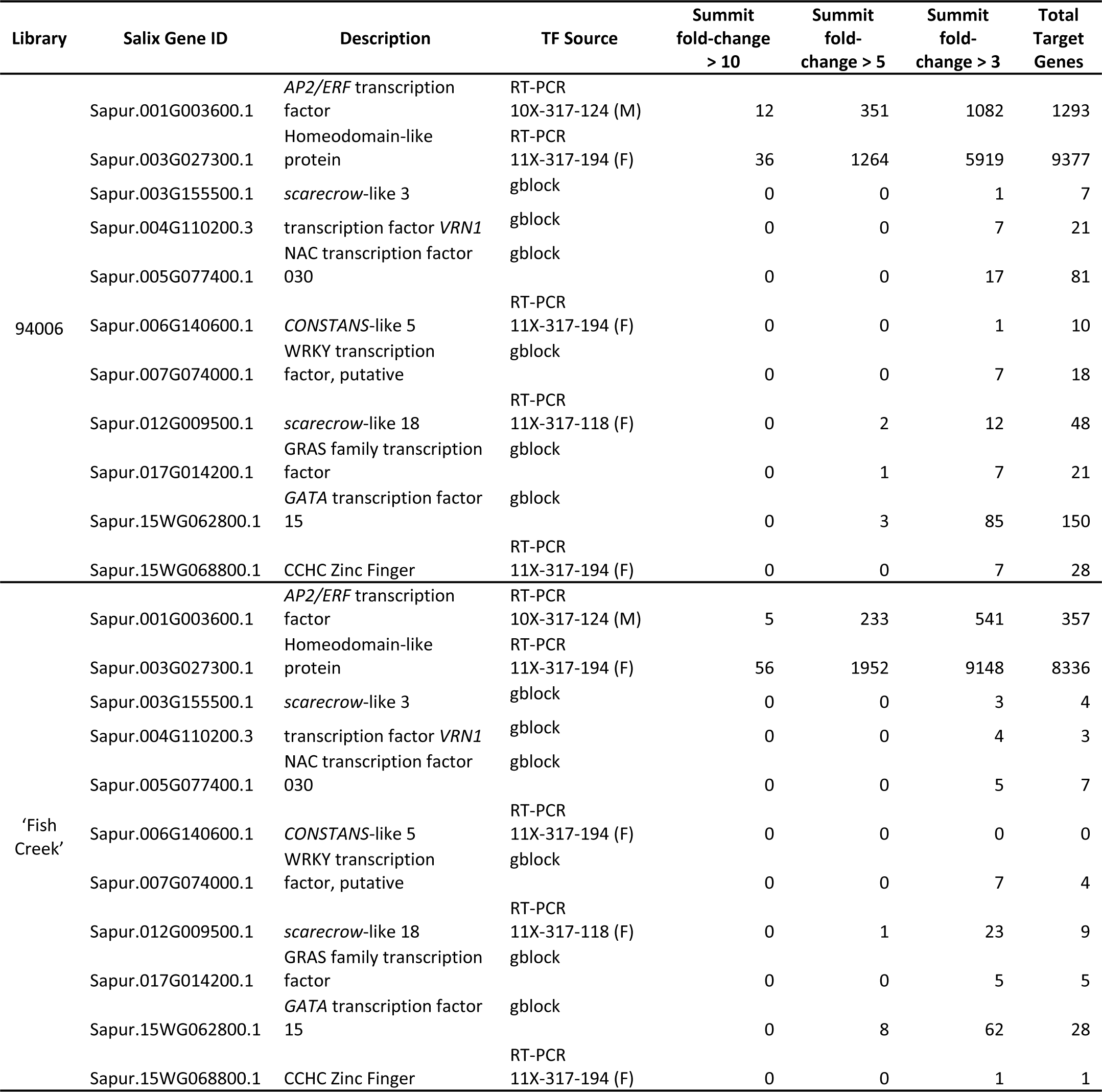
Analyzed transcription factors and libraries with total number of significant summits and target genes from DAP-Seq analysis.

| Library | Salix Gene ID | Description | TF Source | Summit fold-change > 10 | Summit fold-change > 5 | Summit fold-change > 3 | Total Target Genes |
| --- | --- | --- | --- | --- | --- | --- | --- |
| 94006 | Sapur.001G003600.1 | *AP2/ERF* transcription factor | RT-PCR<br>10X-317-124 (M) | 12 | 351 | 1082 | 1293 |
|  | Sapur.003G027300.1 | Homeodomain-like protein | RT-PCR<br>11X-317-194 (F) | 36 | 1264 | 5919 | 9377 |
|  | Sapur.003G155500.1 | *scarecrow*-like 3 | gblock | 0 | 0 | 1 | 7 |
|  | Sapur.004G110200.3 | transcription factor *VRN1* | gblock | 0 | 0 | 7 | 21 |
|  | Sapur.005G077400.1 | NAC transcription factor 030 | gblock | 0 | 0 | 17 | 81 |
|  | Sapur.006G140600.1 | *CONSTANS*-like 5 | RT-PCR<br>11X-317-194 (F) | 0 | 0 | 1 | 10 |
|  | Sapur.007G074000.1 | WRKY transcription factor, putative | gblock | 0 | 0 | 7 | 18 |
|  | Sapur.012G009500.1 | *scarecrow*-like 18 | RT-PCR<br>11X-317-118 (F) | 0 | 2 | 12 | 48 |
|  | Sapur.017G014200.1 | GRAS family transcription factor | gblock | 0 | 1 | 7 | 21 |
|  | Sapur.15WG062800.1 | *GATA* transcription factor 15 | gblock | 0 | 3 | 85 | 150 |
|  | Sapur.15WG068800.1 | CCHC Zinc Finger | RT-PCR<br>11X-317-194 (F) | 0 | 0 | 7 | 28 |
| 'Fish Creek' | Sapur.001G003600.1 | *AP2/ERF* transcription factor | RT-PCR<br>10X-317-124 (M) | 5 | 233 | 541 | 357 |
|  | Sapur.003G027300.1 | Homeodomain-like protein | RT-PCR<br>11X-317-194 (F) | 56 | 1952 | 9148 | 8336 |
|  | Sapur.003G155500.1 | *scarecrow*-like 3 | gblock | 0 | 0 | 3 | 4 |
|  | Sapur.004G110200.3 | transcription factor *VRN1* | gblock | 0 | 0 | 4 | 3 |
|  | Sapur.005G077400.1 | NAC transcription factor 030 | gblock | 0 | 0 | 5 | 7 |
|  | Sapur.006G140600.1 | *CONSTANS*-like 5 | RT-PCR<br>11X-317-194 (F) | 0 | 0 | 0 | 0 |
|  | Sapur.007G074000.1 | WRKY transcription factor, putative | gblock | 0 | 0 | 7 | 4 |
|  | Sapur.012G009500.1 | *scarecrow*-like 18 | RT-PCR<br>11X-317-118 (F) | 0 | 1 | 23 | 9 |
|  | Sapur.017G014200.1 | GRAS family transcription factor | gblock | 0 | 0 | 5 | 5 |
|  | Sapur.15WG062800.1 | *GATA* transcription factor 15 | gblock | 0 | 8 | 62 | 28 |
|  | Sapur.15WG068800.1 | CCHC Zinc Finger | RT-PCR<br>11X-317-194 (F) | 0 | 0 | 1 | 1 |

## Discussion

### Transgenic Arabidopsis and proteomic analysis

In this study we were able to measure and identify over 17,000 total proteins, including 11,000 protein models for each overexpressed gene and a multitude of PDAs (Table 1, Fig. S1-S3). These protein numbers exceed that of recent studies on Arabidopsis floral tissue, which identified between 8,000 and 12,000 proteins (Jing et al., 2020; Lu et al., 2020). This is the first study reporting on *S. purpurea* gene heterologous expression in Arabidopsis, confirming the validity of this system for functional genomics studies in *Salix*, which is especially useful since stable transformants in *S. purpurea* cannot be generated.

Three of the candidate genes used in this study have vague annotations and have not been previously well characterized: Sapur.15WG068800 (pBH102, CCHC Zinc Finger nuclease), Sapur.15WG074900 (pBH104, hypothetical protein), and Sapur.15WG075700 (pBH106, hypothetical protein). The MapMan functional enrichment analysis of PDAs from these genes’ expression lines can provide some insight into their potential role. For Sapur.15WG068800, enriched terms included phosphorylation, carrier-mediated transport, and solute transport channels, which suggest a role in regulating transmembrane transport. In the lines with expression of Sapur.15WG074900, the exceptional number of PDAs observed (5,318) is particularly interesting. Sapur.15WG074900 is a hypothetical protein that unique to Chr15W and female *S. purpurea* and shows high levels of RNA expression in female catkins, but lacks a homolog in Arabidopsis, despite lacking an Arabidopsis homolog. The closest homologs of this gene in *P. trichocarpa* and *P. deltoides* and are also uncharacterized (Tuskan et al., 2006; Goodstein et al., 2011; Hyden et al., 2021). Nevertheless, the large number of floral PDAs suggest that this gene likely has conserved patterns of transcriptional activation in Arabidopsis. The MapMan enrichment categories from the proteomic data suggest a potential role in either directly or indirectly regulating RNA or protein modification and stability. Sapur.15WG075700 is another gene annotated as a hypothetical protein and appears to be unique to *Salix*, as there are no homologs in either *Arabidopsis* or *Populus* (Tuskan et al., 2006; Lamesch et al., 2012). Among the MapMan enriched terms for Sapur.15WG075700 PDAs were microfilament network, primary active transport, and fatty acid metabolism, which together could suggest a role in intracellular transport.

The *ARR17*-OX and *GATA15*-OX lines stood out in this study as having an exceptional number of PDAs when compared with most of the other lines, as well as unique differential expression of multiple proteins with floral development annotations (Tables 1-2, Fig. 2). The downregulation of *PISTILLATA* in the *ARR17* expression lines is particularly interesting. *PISTILLATA* is a well-characterized B-class MADS box gene that is necessary for stamen development (Krizek and Meyerowitz, 1996), which in *S. purpurea* has also been confirmed to have exceptionally high expression in males (Hyden et al., 2021). *ARR17* is hypothesized to act as a switch from male to female development in *Populus* species through the downregulation of *PISTILLATA* expression (Cronk and Müller, 2020). The results from this study support a similar mechanism in *S. purpurea*. Downregulation of *PISTILLATA*, and multiple tapetum and pollen development genes identified exclusively in the *ARR17*-OX lines, along with the differential expression of over 4,300 floral proteins, provides the first functional evidence supporting a role of *ARR17* as a master regulator of sex determination in *S. purpurea*, functioning in the suppression of male floral development (Fig. 3).

**Figure 3.**
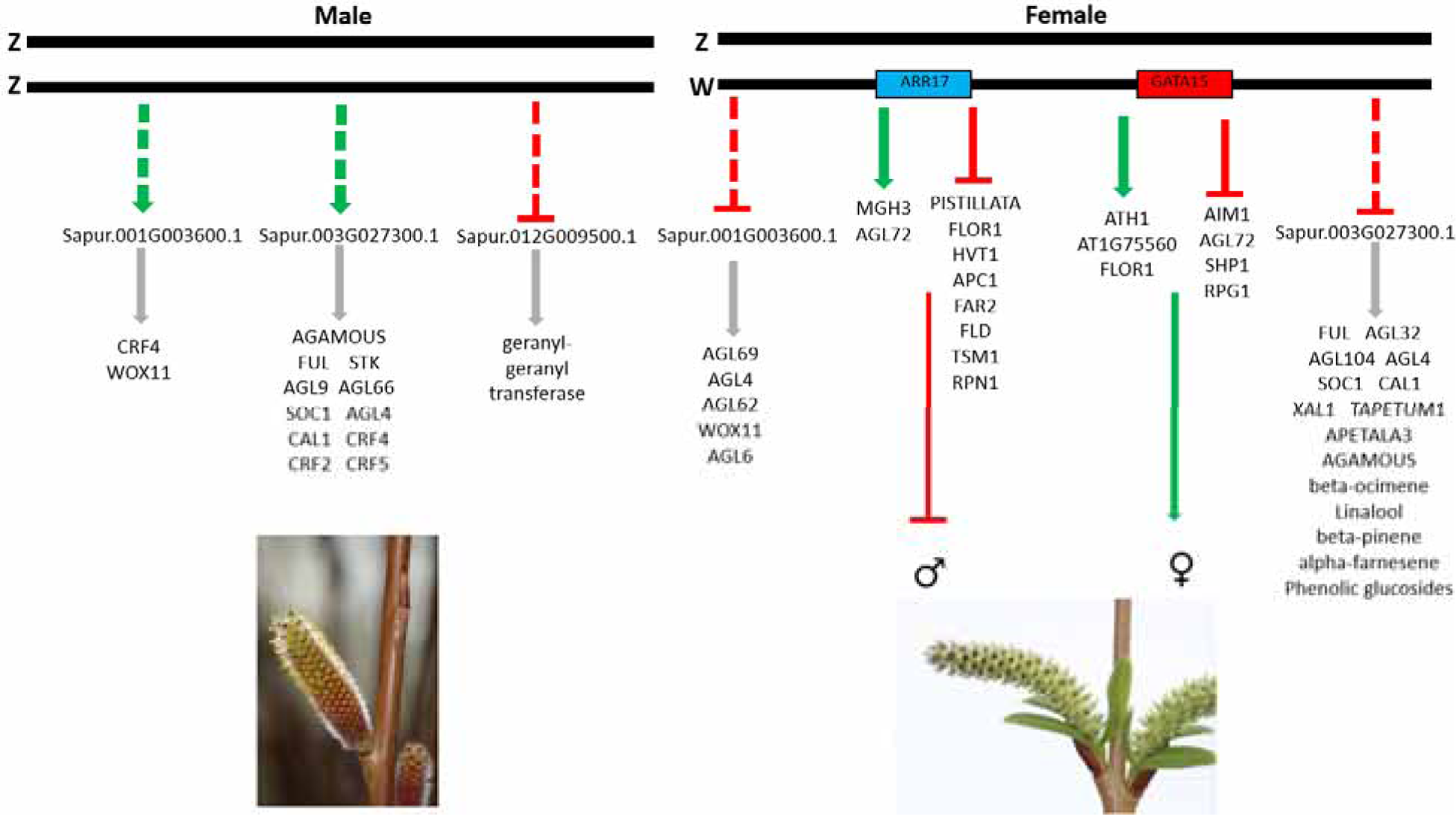
Model for regulation of sex determination in *S. purpurea.* Males represent the default sex when the W chromosome is absent. Genes predicted to be up (green arrow) and down (red line) regulated by the putative master regulators *ARR17* and *GATA15* are shown, which result in simultaneous suppression of male floral development and promotion of female floral development, respectively. Predicted targets of each DAP-Seq transcription factor in males and females that have likely involvement in sex dimorphism are listed, although the exact regulation of expression of each DAP-Seq gene is unknown.

*GATA15* is a proposed master regulator gene of sex determination in *S. purpurea* that shows female-specific expression in mature catkins as well as differential expression in females in early floral shoot development (Carlson et al., 2017; Hyden et al., 2021). It is also the only candidate sex determination gene that is still present on Chr15W in monoecious willows, as other putative master regulator genes were deleted. These monoecious genotypes contain a Chr15W with structural variation and produce both male and female flowers. The presence of *GATA15* and *ARR17* has led us to hypothesize that the role of *GATA15* is to promote female development (Hyden et al., 2023). *GATA15* homologs have been shown to have a role in floral development in Arabidopsis and *Lagerstroemia speciosa* (Ranftl et al., 2016; Hu et al., 2019). In the present study, expression of *S. purpurea GATA15* produced the greatest number of floral PDAs, including several with annotations related to floral development and transition, consistent with previous data that show *GATA15* is expressed early in the transition from vegetative to floral meristem identity when catkin development is determined (Zhang and Fernando, 2005; Carlson et al., 2017). Moreover, among the most downregulated genes in the *GATA15* lines were two isoforms of *LEUNIG*, which is involved in gynoecium development and whose knockout in Arabidopsis has been shown to convert sepals to carpels, reduce stamen number, and alter expression on *PISTILATTA, AGAMOUS*, *AP3*, and *AP1* MADS-box genes (Liu and Meyerowitz, 1995; Lamesch et al., 2012). These proteomic results further support the hypothesized role of *GATA15* as a master regulator of sex determination in *S. purpurea* with an involvement in female floral development.

Taken together, the floral proteomic data from this study indicate that of the genes tested *GATA15* and *ARR17* are the most likely to be master regulators of sex determination in *S. purpurea*, with *ARR17* likely suppressing male flower development and *GATA15* promoting female flower development (Fig. 3). Such a system is consistent with the two-gene model of sex determination, which is common among angiosperms (Charlesworth, 2002) and has been identified in other species, including garden asparagus (Harkess et al., 2020) and kiwifruit (Akagi et al., 2019).

### DAP-Seq

Of the three transcription factors (TFs) that produced consistent binding motifs and large number of significant peaks, two (Sapur.001G003600.1 and Sapur.003G027300.1) have eQTL that map to the SDR and are predicted to be genes in the sex dimorphism pathway that are regulated either directly or indirectly by the master regulator genes of sex (Hyden et al., 2021), while Sapur.15WG062800.1 (*GATA15*) is a candidate master regulator gene. Among the genes targeted by the *AP2/ERF* TF Sapur.001G003600.1 were a *Wuschel*-like *WOX11* in both libraries, a cytokinin response factor 4 in the male ‘Fish Creek’, and four MADS-box genes: *AGL4*, *AGL6*, *AGL62*, and *AGL69*, in the female 94006. The *AGL4* target is particularly interesting, as it is involved in ovule development (Rounsley et al., 1995), and showed a 4.96-fold increase in TF binding in the 94006 library. This preponderance of MADS-box gene binding exclusively in the female library suggests a potential role of this TF in promoting female floral development.

Sapur.003G027300.1 exhibited binding near both floral development genes and secondary metabolism genes. Among the floral development genes with adjacent TF binding in both libraries were homologs of two *AGAMOUS*-like genes (involved stamen and ovule identity) (Mizukami and Ma, 1992), *AGL32* (ovule endothelial identity) (De Folter et al., 2006), and *AGL4* (ovule development) (Rounsley et al., 1995), pointing towards involvement of this gene in floral development in both sexes. In the 94006 library, Sapur.003G027300.1 binding sites were identified observed near *TAPETUM1*, *TPD1*, and *AP3* (Krizek and Meyerowitz, 1996; Yang et al., 2003; Lamesch et al., 2012), all of which are directly involved in stamen development and may suggest a role of this gene in downregulating male floral development in females. Furthermore, Sapur.003G027300.1 displayed binding near multiple cytokinin response factor genes exclusively in the male ‘Fish Creek’ library, indicating a role in regulating cytokinin in males. This is particularly interesting considering the proposed role of *ARR17*, a cytokinin response regulator, in sex determination in *S. purpurea* (Zhou et al., 2020; Hyden et al., 2021). Indeed, given that eQTL for this gene map to the sex determination region (Hyden et al., 2021), it is possible that *ARR17* may directly or indirectly regulate expression of Sapur.003G027300.1, although the precise mechanism for this remains unclear. Sapur.003G027300.1 also binds near a multitude of genes involved in terpenoid, phenolic glucoside, and flavonoid production in the 94006 library, including genes specifically annotated as being involved in production of beta-ocimene, beta-pinene, limonene, and alpha-farnesene. These aforementioned compounds are terpenoids that are differentially produced in male and female *S. purpurea* catkins and are associated with pollinator and pest attraction (Keefover-Ring et al., 2022). TF binding activity near genes responsible for production of these metabolites in 94006 but not ‘Fish Creek’ suggests that, under the presence of the Chr15W and the sex determination genes, the CpG methylation profile is altered, which in turn affects TF binding and regulation of these metabolites in females resulting in differential expression and sex dimorphism.

The DAP-Seq assay of Sapur.15WG062800 *GATA15*, a candidate master regulator gene, showed peaks associated with *ATH1* and a CCHC Zinc finger in 94006. *ATH1* is activated by the C class MADS box gene *AGAMOUS*, and in turn regulates GA synthesis (Gómez-Mena et al., 2005). The CCHC Zinc finger targeted by *GATA15* is also described as having a likely role in reproductive development in Arabidopsis (Lamesch et al., 2012). This binding activity of Sapur.15WG062800 was only observed in the female 94006 library. These data are consistent with previous research hypothesizing *GATA15* as a female-specific master regulator gene of sex in *S. purpurea* with a role in promoting female floral development (Hyden et al., 2023).

The targeting of different gene families involved in sex dimorphism between male and female libraries was consistently observed across three transcription factors tested with DAP-Seq and further supports a role for CpG methylation, which is nearly three-fold higher in male *S. purpurea* catkins compared to females, in sex dimorphism (Hyden et al., 2021).

### Comparison of Arabidopsis heterologous expression and DAP-Seq results

Among the *Salix* genes tested in this study, the CCHC zinc finger Sapur.15WG098800 gene and *GATA15* Sapur.15WG062800 gene were both analyzed through heterologous expression in Arabidopsis and in DAP-Seq assays, due to their annotation as transcription factors and hypothesized role as master regulators of sex. In both cases, the DAP-Seq and proteomics results provided complementary data supporting or rejecting a role in sex determination. For the CCHC zinc finger Sapur.15WG068800 gene, the fewest PDAs were observed (103) out of any of the transgenic Arabidopsis lines, and the DAP-Seq results indicated only seven and one significant peaks in the female and male libraries, respectively.

Moreover, none of these PDAs or genes adjacent to DAP-Seq peaks had annotations that would suggest a role in regulating sex determination. From these data, the precise function of Sapur.15WG06800 remains inconclusive, but a role in regulating sex determination seems unlikely. For *GATA15* on the other hand, both the DAP-Seq and the Arabidopsis expression data supported a potential role in sex determination as a promoter of female development, with the greatest number of PDAs in the Arabidopsis heterologous expression lines, multiple floral development proteins with unique abundance patterns, and DAP-Seq peaks near two well-characterized floral development genes.

In summary, results from this study support the role of *ARR17* and *GATA15* as master regulator genes of sex determination in *S. purpurea*. *ARR17* appears to suppress expression of *PISTILLATA* and tapetum development genes, implicating a role as a male suppressor gene, while *GATA15* appears to promote female floral development by regulating expression of floral transition and ovule development genes, including *LEUNIG*. This system is clearly distinct from the single-gene system in *Populus* and underscores the dynamic nature of the sex determination system in the Salicaceae family.

## Materials and Methods

### Generation and evaluation of Arabidopsis heterologous expression lines

*Salix purpurea* genes used for heterologous expression in Arabidopsis were obtained from the list of eight candidate sex-determination genes described in a previous study (Hyden et al., 2021) including Arabidopsis Response Regulator 17 (*ARR17*) (Sapur.15WG073500), Double-stranded RNA-Binding 1 (*DRB1*) (Sapur.15WG074300), a CCHC znc finger nuclease (Sapur.15WG068800), *GATA15* (Sapur.15WG062800), and two genes annotated as hypothetical proteins (Sapur.15WG074900, Sapur.15WG075700). One gene, Sapur.15WG074400, a homolog of *AGO4*, did not contain either a start codon or a canonical stop codon, and was therefore dropped from further consideration. Attempts to clone Sapur15WG075300, annotated as a hypothetical protein, did not produce any colonies containing the transgene in *E. coli*, suggesting that it may result in a toxic product, and therefore this gene was also dropped from further consideration. *Salix purpurea* homologs of *Flowering Locus T* (*FT*, Sapur.008G061900) and *LEAFY* (Sapur.15WG122200) were included as positive controls to test the effectiveness of the transformation methods and validity of the results, since their role and function are well characterized in Arabidopsis. All coding sequences (CDS) were obtained from the *S. purpurea* female 94006 v5.1 reference (Zhou et al., 2020) available on Phytozome (Goodstein et al., 2011). CDSs were synthesized as gblocks by Integrated DNA Technologies (Coralville, IA, USA) and contained 30 bp and 26 bp overlap sequences homologous to the pGFPGUSPlus vector (Vickers et al., 2007) on the 3’ and 5’ ends, respectively, along with a six His tail immediately prior to the stop codon. NEBuilder HiFi DNA assembly (New England Biolabs, Ipswich, MA, USA) was used to assemble each gblock into the pGFPGUSPlus vector, replacing the GFP CDS adjacent to a 35S promoter (Supplemental Datasets S1-S8). Each vector contained plant selectable markers for GUS and hygromycin, also driven by 35S promoters, as well as a bacterial selectable marker for kanamycin resistance. Assembled constructs were used to transform chemically competent TOP10 *E. coli* cells obtained from ThermoFisher (Waltham, MA, USA) following the manufacturer’s protocol. Successful insertion of each gblock in the correct orientation was confirmed by restriction digest and PCR amplification using Q5 polymerase (New England Biolabs, Ipswich, MA, USA) followed by Sanger sequencing of plasmid DNA in the Cornell University Institute for Biotechnology (Ithaca, NY, USA). Plasmid DNA was extracted from *E. coli* cells using a miniprep kit from Qiagen (Germantown, MD, USA) following the recommended protocol. *Agrobacterium tumefaciens* GV3101 cells were transformed using a standard electroporation protocol, and insertion of each plasmid and gene of interest were confirmed with PCR using Q5 polymerase followed by Sanger sequencing.

Columbia-0 ecotype Arabidopsis was transformed by floral dip following the protocol described by (Zhang et al., 2006). T_1_ generation Arabidopsis seeds were grown on standard MS media containing hygromycin for selection of transformants at 30 µg L^-1^ and surviving seedlings were transferred to potting mix. The presence of each transgene was confirmed using PCR from genomic DNA followed by Sanger sequencing of PCR products. Four to six T_1_ plants, each representing a unique transgene insertion event, were self-pollinated to generate T_2_ seeds, which were grown on MS media containing 30 µg L^-1^ hygromycin before being transferred to potting mix. All Arabidopsis were grown at 21°C under fluorescent lighting with an 8/16 hour photoperiod prior to bolting. Upon initiation of floral shoots, day length was switched to 16 hours. Plants were sub-irrigated regularly whenever the potting mix became dry. Expression of the plasmid in floral tissue of T_2_ plants was confirmed by GUS staining assay following the manufacturer’s protocol (Millipore Sigma, Burlington, MA, USA) (Figure S6). Floral buds from three T_2_ plants (biological replicates) with confirmed gene insertion events for each event were harvested prior to anthesis, flash frozen in liquid nitrogen, and stored at -80°C.

### Protein Extraction and Proteome Analysis

Three floral buds collected prior to anthesis from each from three biological replicates (full sibling T_2_ plants) for each transgenic insertion event were selected for proteomics. Floral buds were ground in liquid nitrogen using 2.3mm zirconia/silica beads with a Geno/Grinder 2010 (SPEX) at a rate of 1200rpm for 1 minute. Ground tissue was resuspended in 200µL of lysis buffer (4% sodium dodecyl sulfate, 10mM dithiothreitol) and incubated for 10 minutes at 90°C with constant shaking. Proteins were alkylated with 30mM iodoacetamide and incubated in the dark for 15 minutes to prevent the reformation of disulfide bonds. For each sample, all of the crude protein extract was transferred to a fresh tube, Sera-Mag beads were added (100µg), and proteins were extracted by protein aggregation capture (Batth et al., 2019).

Precipitated protein was resuspended in 100mM ammonium bicarbonate (ABC) and then digested with two separate and sequential aliquots of sequencing grade trypsin (Promega) in the ratio of 1:75 trypsin to sample protein ratio overnight followed by a 3-hr digestion. Peptide mixtures were adjusted to 0.5% formic acid (FA) and physically separated from the Sera-Mag beads with an AcroPrep Advance 96-well 10KDa omega filter plate (Pall Corporation) by centrifuging at 1500xg for 30 minutes. Peptides were freeze dried (Labconco FreeZone 72040) and then resuspended in an aqueous solvent (0.1% formic acid, 5% ACN). Peptide concentrations were estimated using a Nanodrop One spectrophotometer. For each sample, 2µg aliquots were measured by one-dimensional liquid chromatography tandem mass spectrometry (1D-LC-MS/MS) using a RSLCnano UHPLC system (Thermo Scientific) coupled to a Q Exactive Plus mass spectrometer (Thermo Scientific). Peptide mixtures were first injected across an in-house built strong cation exchange (SCX) Luna trap column (5 μm, 150 μm X 50mm; Phenomenex, USA) followed by a nanoEase symmetry reverse phase (RP) C18 trap column (5 μm, 300 μm X 50mm; Waters, USA) and then washed with the aqueous solvent. A 1M ammonium acetate inject was used to elute peptides to the C18 trap column, which was then switched to be in-line with an in-house pulled nanospray emitter analytical column (75 μm X 350 mm) packed with Kinetex RP C18 resin (1.7 μm; Phenomenex, USA). Peptides were separated over a 160-minute linear gradient from 2 to 25% of mobile phase (0.1% FA, 80% ACN) at a flow rate of 250nL/min and analyzed using a Top10 data dependent acquisition strategy (Villalobos Solis et al., 2019). All MS data were acquired with Thermo Xcalibur (version 4.2.47) and analyzed using the Proteome Discoverer software (Thermo-Fisher Scientific, version 2.5) (Orsburn, 2021). Each MS raw data file was processed by the SEQUEST HT database search algorithm (Eng et al., 1994) and confidence in peptide-to-spectrum (PSM) matching was evaluated by Percolator (Käll et al., 2007). The TAIR 11 reference genome (Lamesch et al., 2012) was used for mapping proteins. Peptide and PSMs were considered identified at q < 0.01 and proteins were required to have at least one unique peptide sequence. Proteins with at least one unique peptide were exported from Proteome Discoverer. Log2-transformation of protein abundances was performed followed by local regression (LOESS) normalization and mean-centering across the entire dataset in R using scripts from the InfernoRDN software (v1.1.7995) (Larsson, 2014). The abundance values for proteins with missing values were imputed with random values drawn from the normal distribution (width 0.3, downshift 2.2) using R. Proteins of differential abundance (PDAs) were calculated using the “limma” package in R (Ritchie et al., 2015) by comparing all events and biological replicates for each overexpressed gene against the pGFPGUSPlus empty vector controls. Volcano plots were generated using EnhancedVolcano (v.1.13.2) (Blighe et al., 2018) with a fold change cutoff of 2 and p-value threshold of 0.01. Mapman functional categories (Schwacke et al., 2019) were assigned to each FASTA sequence using the Mercator4 online submission tool (https://www.plabipd.de/portal/web/guest/mercator4). Functional enrichment for MapMan categories was performed for each gene expression relative to the empty vector control using clusterProfiler (v.4.2.2) (Wu et al., 2021). Briefly, each protein was assigned to a MapMan category, significantly differentially enriched proteins (|logFC|>2 and p.val<0.05) were used as the differentially enriched genes, while all genes in each comparison were used as background. Each differentially abundant protein set was searched for genes with annotations that are well characterized in floral development.

### Transcription factor cloning for DAP-Seq

Genes for DAP-Seq analysis were selected from among 97 transcription factors (TFs) identified as having a potential role as top level regulator genes of sex determination from an eQTL study (Hyden et al., 2021). These genes were prioritized based on transcription factor family, likelihood of success in a DAP-Seq assay, functional annotation, and floral differential gene expression (O’Malley et al., 2016; Hyden et al., 2021). The 11 genes with the highest prioritization score were advanced for DAP-Seq analysis (Table 2). TF CDS regions were successfully cloned via RT-PCR from catkin RNA obtained from the *S. purpurea* 317 F_2_ family (Hyden et al., 2021), or were generated using gblocks from IDT based on the female 94006 v5.1 reference genome (Zhou et al., 2020). Each of the 11 genes were cloned into a pENTR-DTOPO vector and finally into a pIX-HALO expression vector using a Gateway cloning kit obtained from NEB and following the manufacturer’s protocol (Supplemental Datasets S9-S19). PCR followed by Sanger sequencing of plasmid DNA was used to confirm the presence and correct sequence and orientation of each CDS sequence in the entry and destination vectors. Genomic DNA for the DAP-Seq assay was extracted from catkins of female (clone 94006) and male (clone ‘Fish Creek’) *S. purpurea* using a modified Qiagen plant mini kit protocol.

### DAP-Seq experiments

DAP-Seq experiments were conducted as described previously in (O’Malley et al., 2016), with minor modifications, described in (Baumgart et al., 2021). DNA libraries were prepared by fragmenting genomic DNA of either *S. purpurea* ‘Fish Creek’ or *S. purpurea* 94006 using a Covaris LE220-Plus focused-ultrasonicator (Covaris), followed by library preparation with the KAPA HyperPrep kit (Roche) following the manufacturer’s recommendations. Insert sizes were targeted to an average of 150 bp. Before use in the DAP-seq assay, libraries were PCR amplified for 10 cycles.

For *in vitro* protein expression, linear fragments were first PCR amplified from each pIX-HALO plasmid using primers targeting the upstream T7 promoter (5’ GTGAATTGTAATACGACTCACTATAGGG 3’) and downstream of the poly-A stretch (5’ CAAGGGGTTATGCTAGTTATTGCTC 3’). The correct size of each PCR product was verified using a Tapestation (Agilent Technologies), and PCR products were purified using SPRI beads. Transcription factors were expressed using at least 2000 ng PCR product per sample with the TnT T7 Quick for PCR DNA in vitro protein expression kit (Promega). All reaction volumes were doubled to yield a total of 100 µL protein product per transcription factor. Each DAP-Seq reaction was run with 100 µL expressed protein, 150 ng of the previously prepared fragment library, and 15 µg salmon sperm DNA to reduce non-specific binding. The final DAP-seq libraries were pooled for sequencing on a NovaSeq using the S4 flowcell (Illumina), targeting 30 million 2x150 reads per sample. Primary data analyses included quality filtering, alignment to the reference genome, peak-calling, and gene assignment as described in (Baumgart et al., 2021). Binding motifs for transcription factors were predicted using MEME version 5.3.0 (Bailey and Elkan, 1994).

## Acknowledgements

The authors are grateful for excellent technical support provided by Michael Quade, McKenzie Schessl, and Alexander Wares. We appreciate insightful comments on the manuscript from Steven DiFazio, Jeffrey Doyle, and Jocelyn Rose.

## Author Contributions

B.L.H., J.G.C., X.Y., R.L.H., G.A.T., R.O., and L.B.S. designed the research; B.L.H., D.L.C., P.E.A., G.Y., T.Y., L.B., Y.Z., C.C., and R.O. performed the experiments, B.L.H. wrote the paper with contributions from all authors.

## Data Availability

All proteomics spectral data in this study were deposited at the ProteomeXchange Consortium via the MASSIVE repository (https://massive.ucsd.edu/). The data can be reviewed under the username “reviewer_MSV000091180” and password “BHArabidopsis”

## Supplemental Datasets

**Supplemental Dataset S1.** Plasmid map (.dna format) for *ARR17* Sapur.15WG073500 expression plasmid (pBH100)

**Supplemental Dataset S2.** Plasmid map (.dna format) for *GATA15* Sapur.15WG062800 expression plasmid (pBH101)

**Supplemental Dataset S3.** Plasmid map (.dna format) for CCHC Zinc Finger Sapur.15WG068800 expression plasmid (pBH102)

**Supplemental Dataset S4.** Plasmid map (.dna format) for *DRB1* Sapur.15WG074300 expression plasmid (pBH103)

**Supplemental Dataset S5.** Plasmid map (.dna format) for hypothetical protein Sapur.15WG074900 expression plasmid (pBH104)

**Supplemental Dataset S6.** Plasmid map (.dna format) for hypothetical protein Sapur.15WG075700 expression plasmid (pBH106)

**Supplemental Dataset S7.** Plasmid map (.dna format) for *LEAFY* Sapur.15WG122200 expression plasmid (pBH107)

**Supplemental Dataset S8.** Plasmid map (.dna format) for *FT* Sapur.008G061900 expression plasmid (pBH108)

**Supplemental Dataset S9.** Plasmid map (.dna format) for pIX-HALO expression vector with Sapur.15WG062800 (pBH217)

**Supplemental Dataset S10.** Plasmid map (.dna format) for pIX-HALO expression vector with Sapur.15WG068800 (pBH218)

**Supplemental Dataset S11.** Plasmid map (.dna format) for pIX-HALO expression vector with Sapur.012G009500 (pBH219)

**Supplemental Dataset S12.** Plasmid map (.dna format) for pIX-HALO expression vector with Sapur.006G140600 (pBH220)

**Supplemental Dataset S13.** Plasmid map (.dna format) for pIX-HALO expression vector with Sapur.007G074000 (pBH225)

**Supplemental Dataset S14.** Plasmid map (.dna format) for pIX-HALO expression vector with Sapur.005G077400 (pBH226)

**Supplemental Dataset S15.** Plasmid map (.dna format) for pIX-HALO expression vector with Sapur.003G155500 (pBH227)

**Supplemental Dataset S16.** Plasmid map (.dna format) for pIX-HALO expression vector with Sapur.004G110200 (pBH228)

**Supplemental Dataset S17.** Plasmid map (.dna format) for pIX-HALO expression vector with Sapur.017G014200 (pBH229)

**Supplemental Dataset S18.** Plasmid map (.dna format) for pIX-HALO expression vector with Sapur.001G003600 (pBH232)

**Supplemental Dataset S19.** Plasmid map (.dna format) for pIX-HALO expression vector with Sapur.003G027300 (pBH242)

**Supplemental Dataset S20.** Expression data on all significant differentially abundant proteins for Arabidopsis expression lines 100 to 108.

**Supplemental Dataset S21.** Listing of all significant target genes for each DAP-Seq assay.

## Supplemental Figures

**Figure S1.**
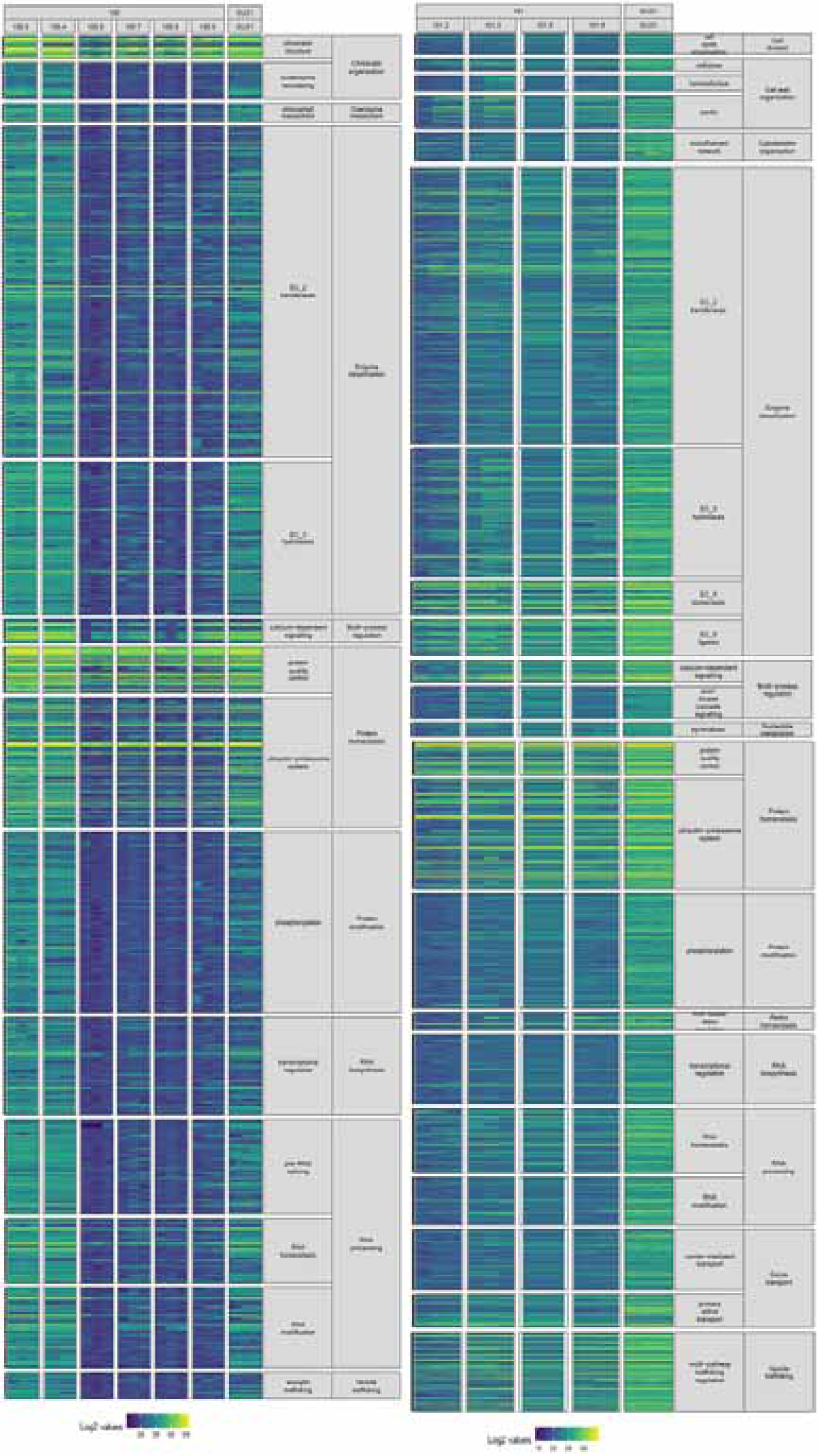
Heatmap displaying the total expression of significantly differentially abundant proteins in the MapMan enriched functional categories for the *ARR17* (100) and *GATA15* (101) expression lines, compared to the pGFPGUSPlus empty vector control (GUS1). Each column represents data from a unique transgene event.

**Figure S2.**
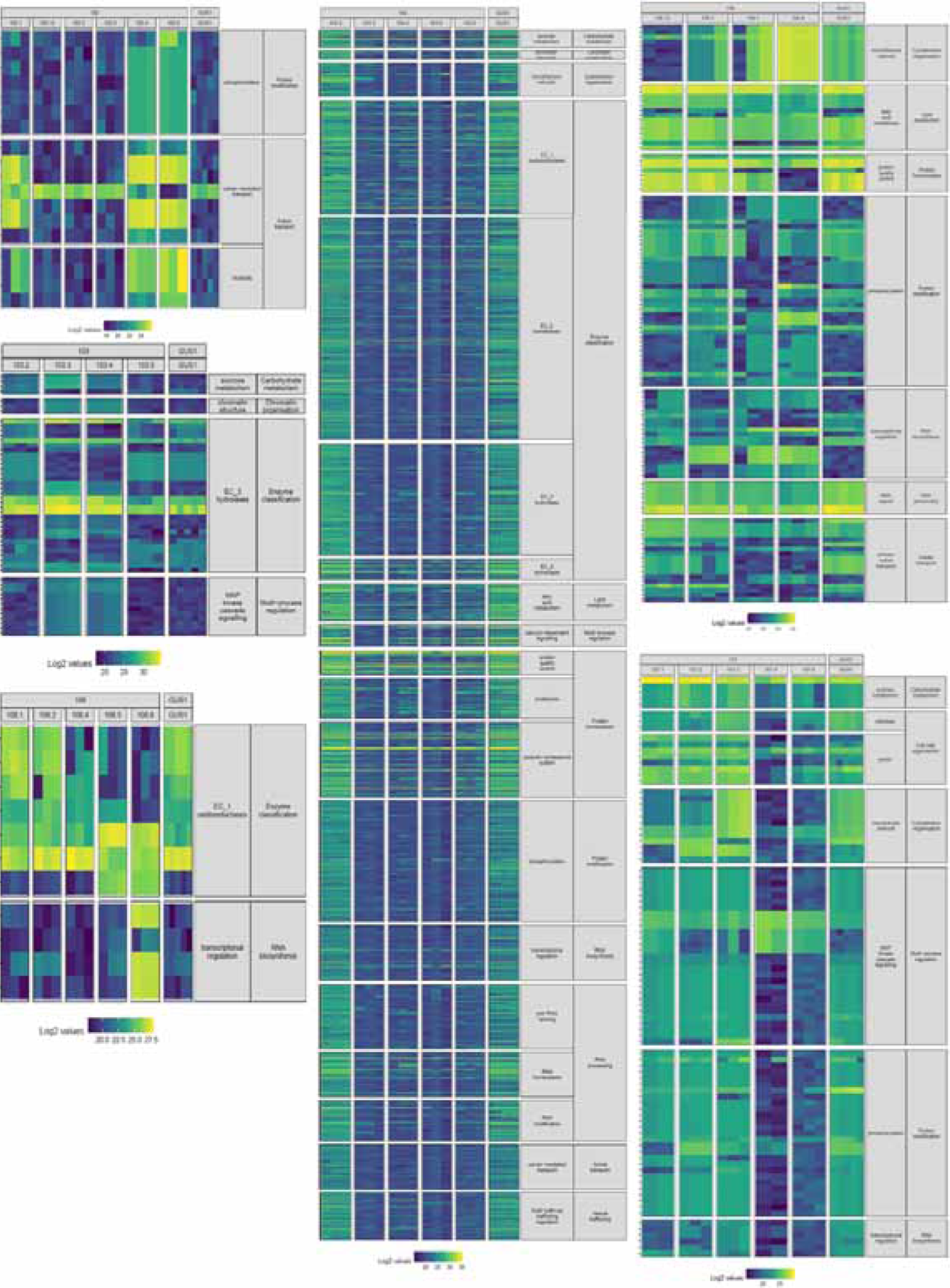
Heatmap displaying the total expression of each gene in the MapMan enriched functional categories for the CCHC Zinc Finger (102) *DRB1* (103) Sapur.15WG074900 hypothetical protein (104), Sapur.15WG075700 hypothetical protein (106), *LEAFY* (107), and *FT* (108) expression lines, compared to the pGFPGUSPlus empty vector control (GUS1). Each column represents data from a unique transgene event.

**Figure S3.**
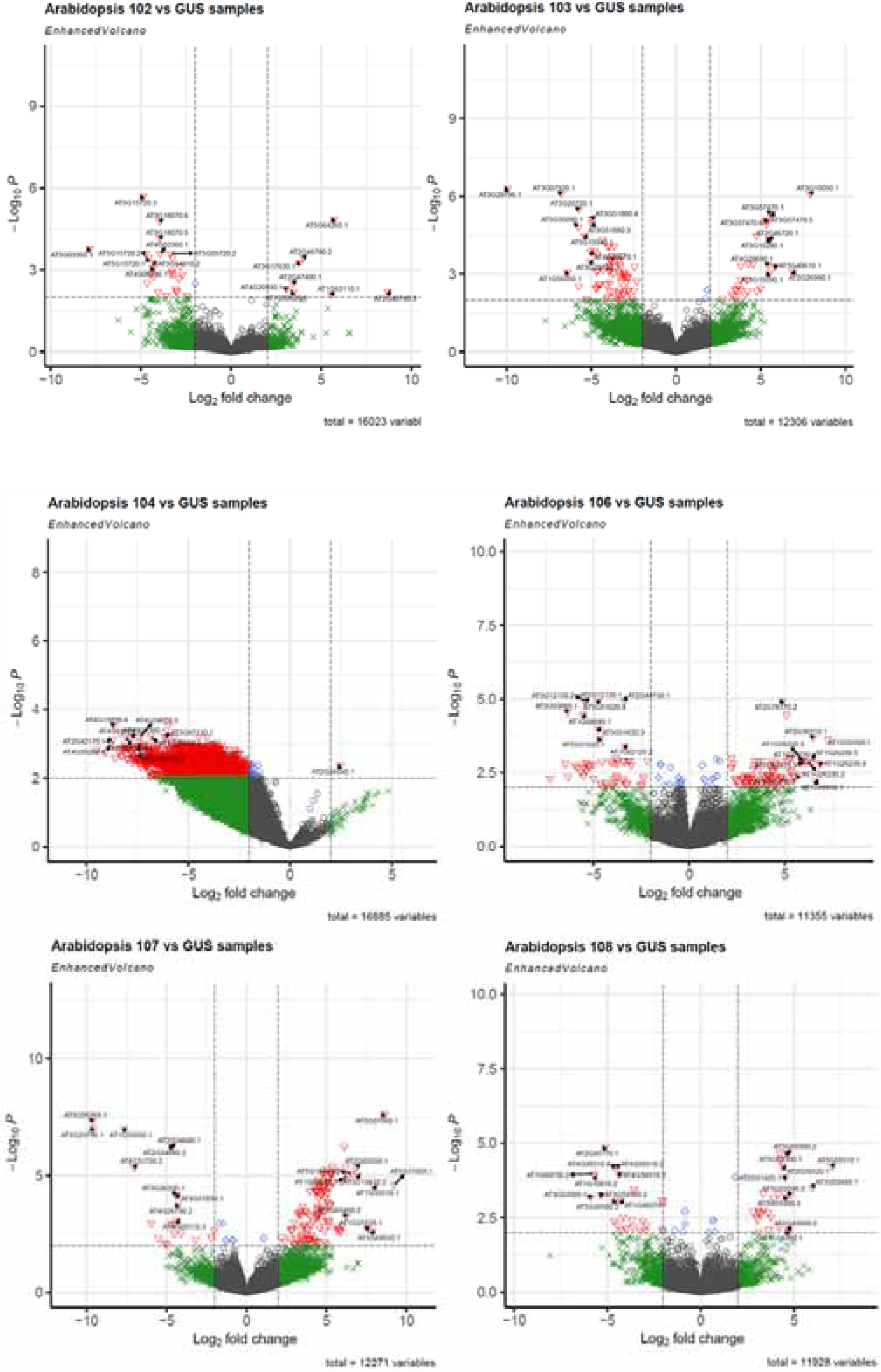
Volcano plots displaying the total differential abundant protein results and top ten significant up and down regulated proteins for the Sapur.15WG068800 CCHC Zinc Finger nuclease (102), Sapur.15WG074300 *DRB1* (103), Sapur.15WG074900 hypothetical protein (104), Sapur.15WG075700 hypothetical protein (106), *LEAFY* (107), and *FT* (108) expression lines, relative to the empty vector control.

**Figure S4.**
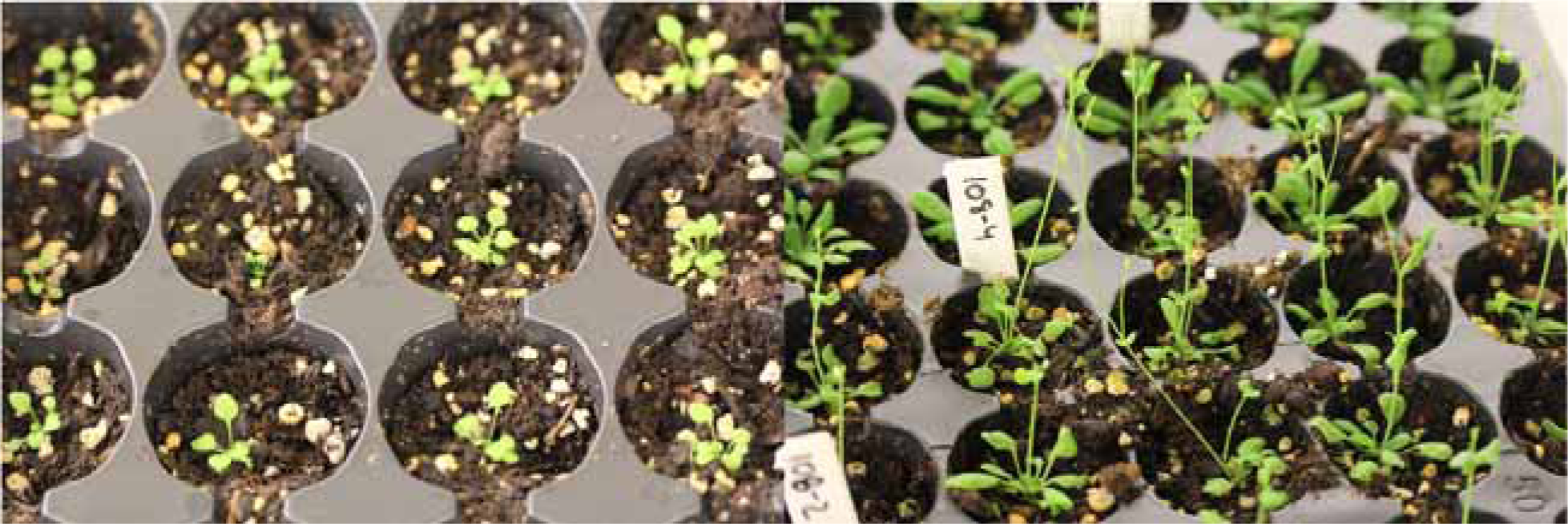
Comparison of Arabidopsis transformed with the empty vector control (left) and overexpressing *S. purpurea FT* (Sapur.008G061900, right)

**Figure S5.**
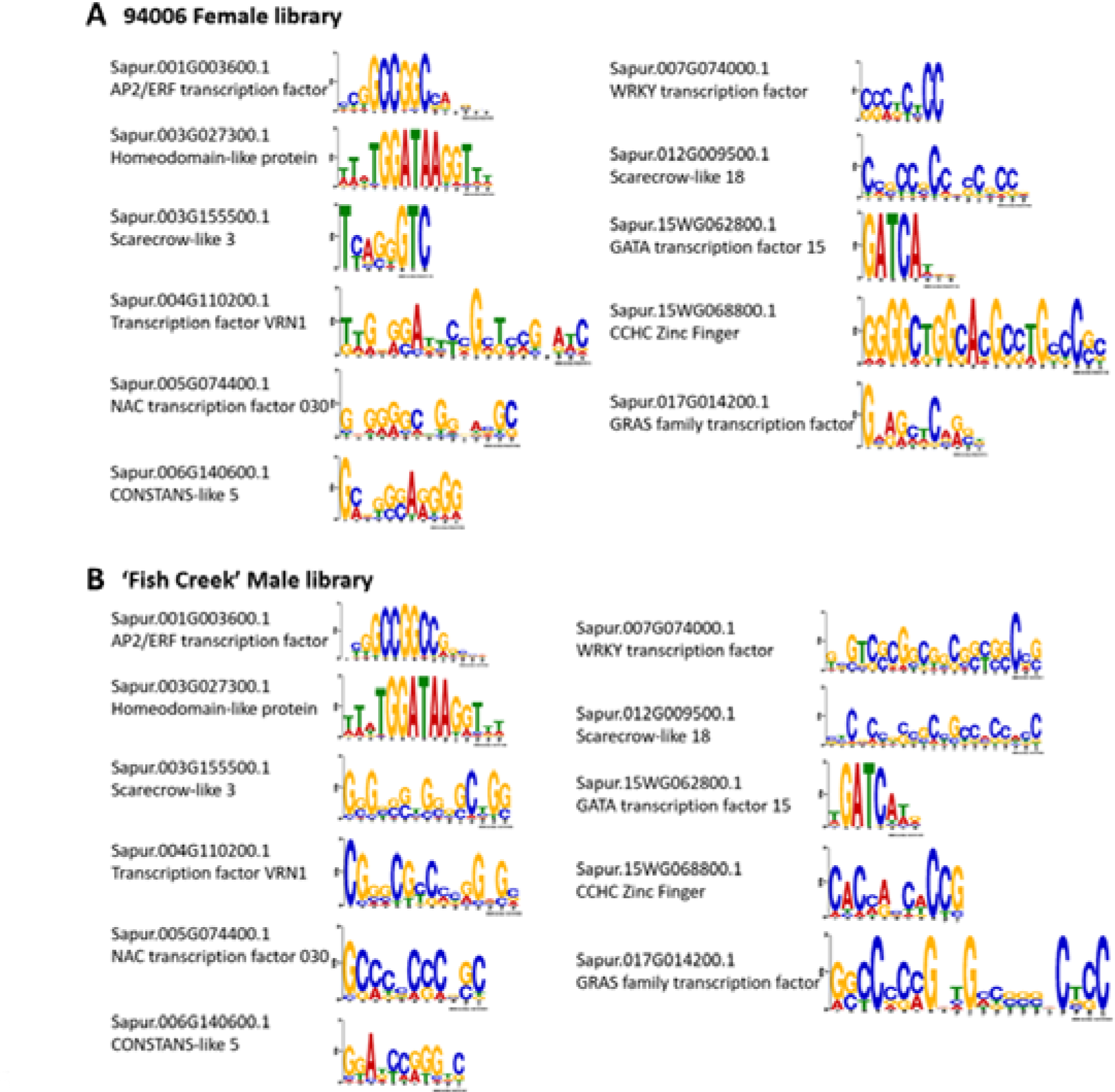
Predicted binding motifs for each transcription factor tested in DAP-Seq. A. Predictions from the 94006 library (female); B. Predictions from the ‘Fish Creek’ library (male).

**Figure S6.**
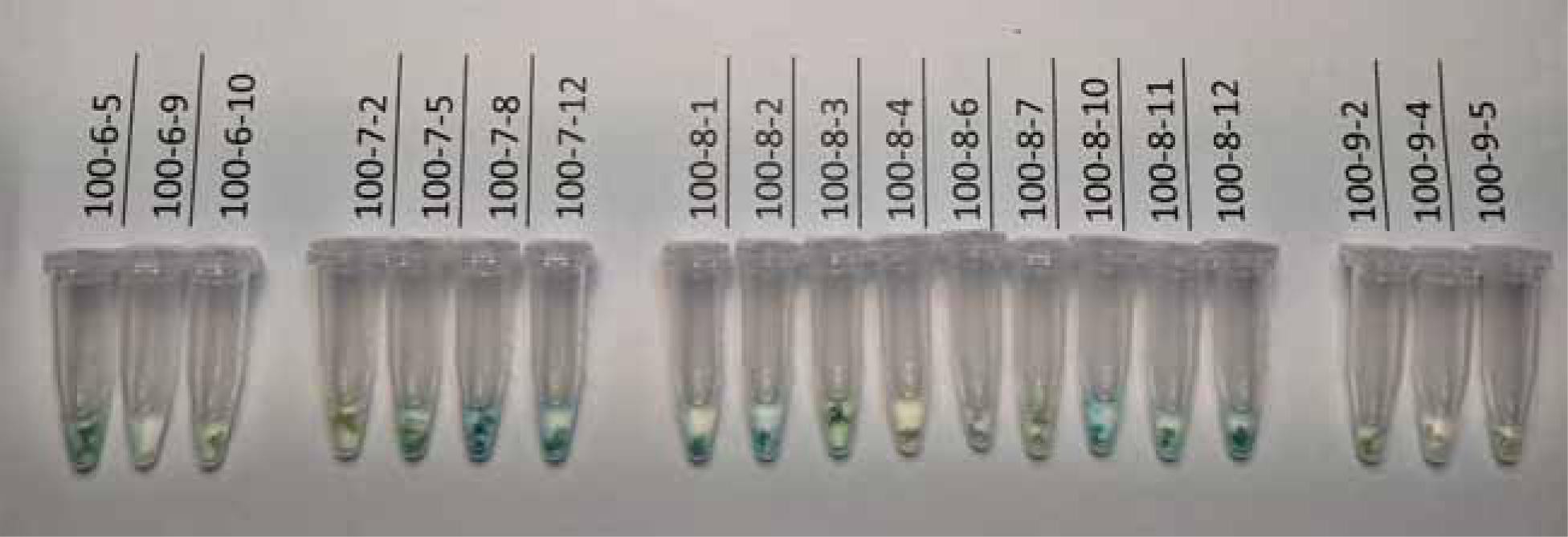
GUS staining assay results for representative samples of T_2_ flowers after ethanol staining

**Table S1.**
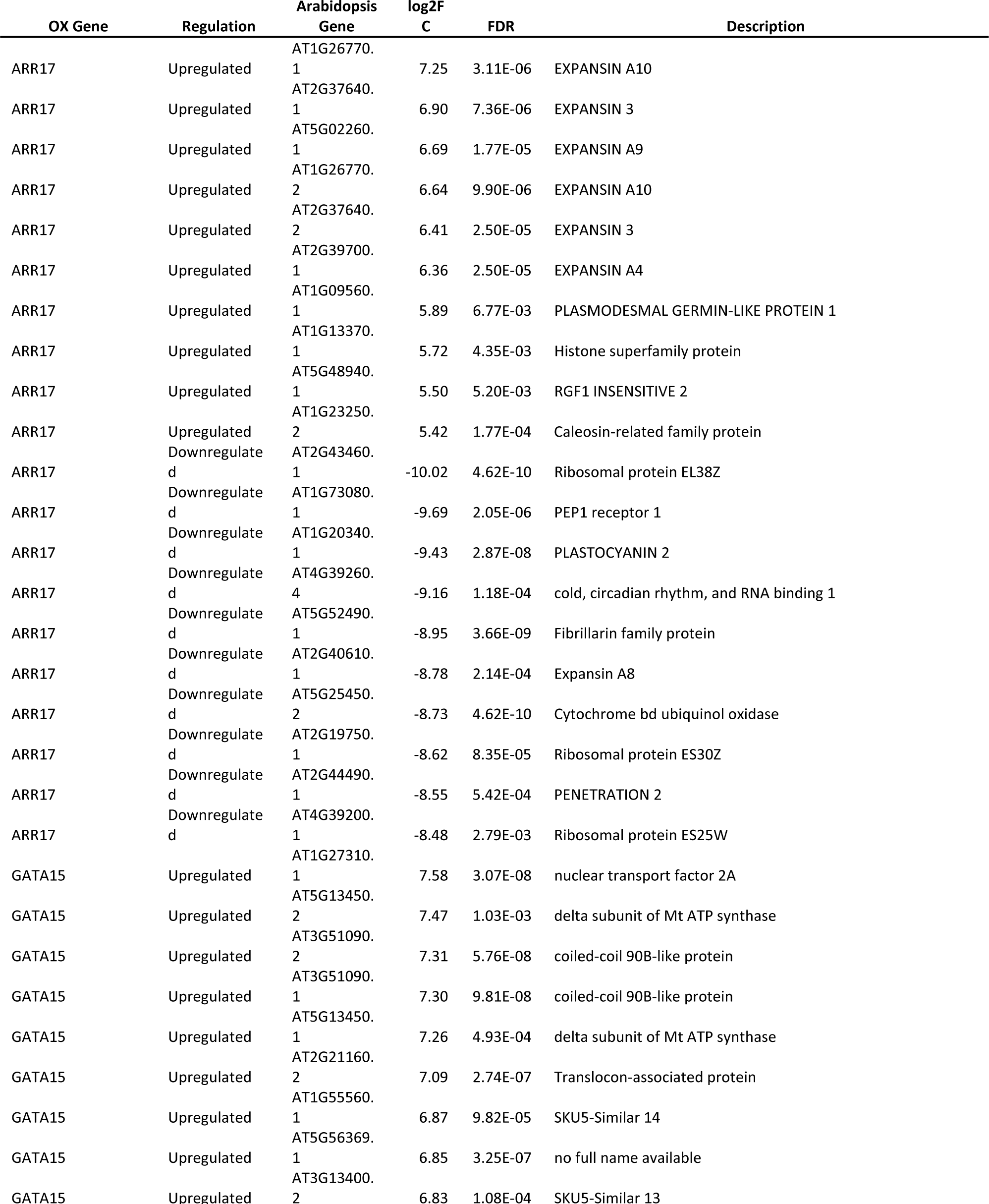
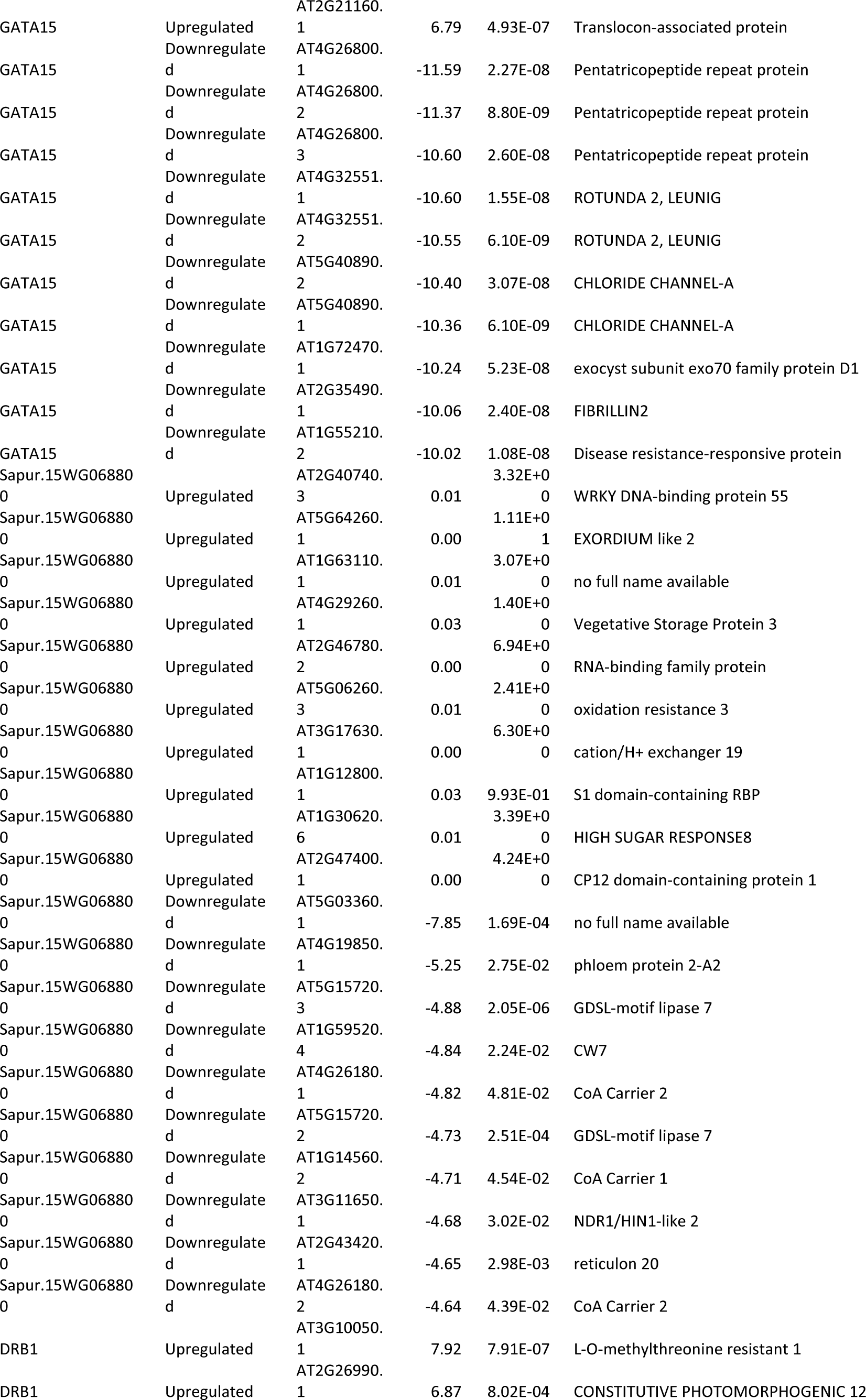
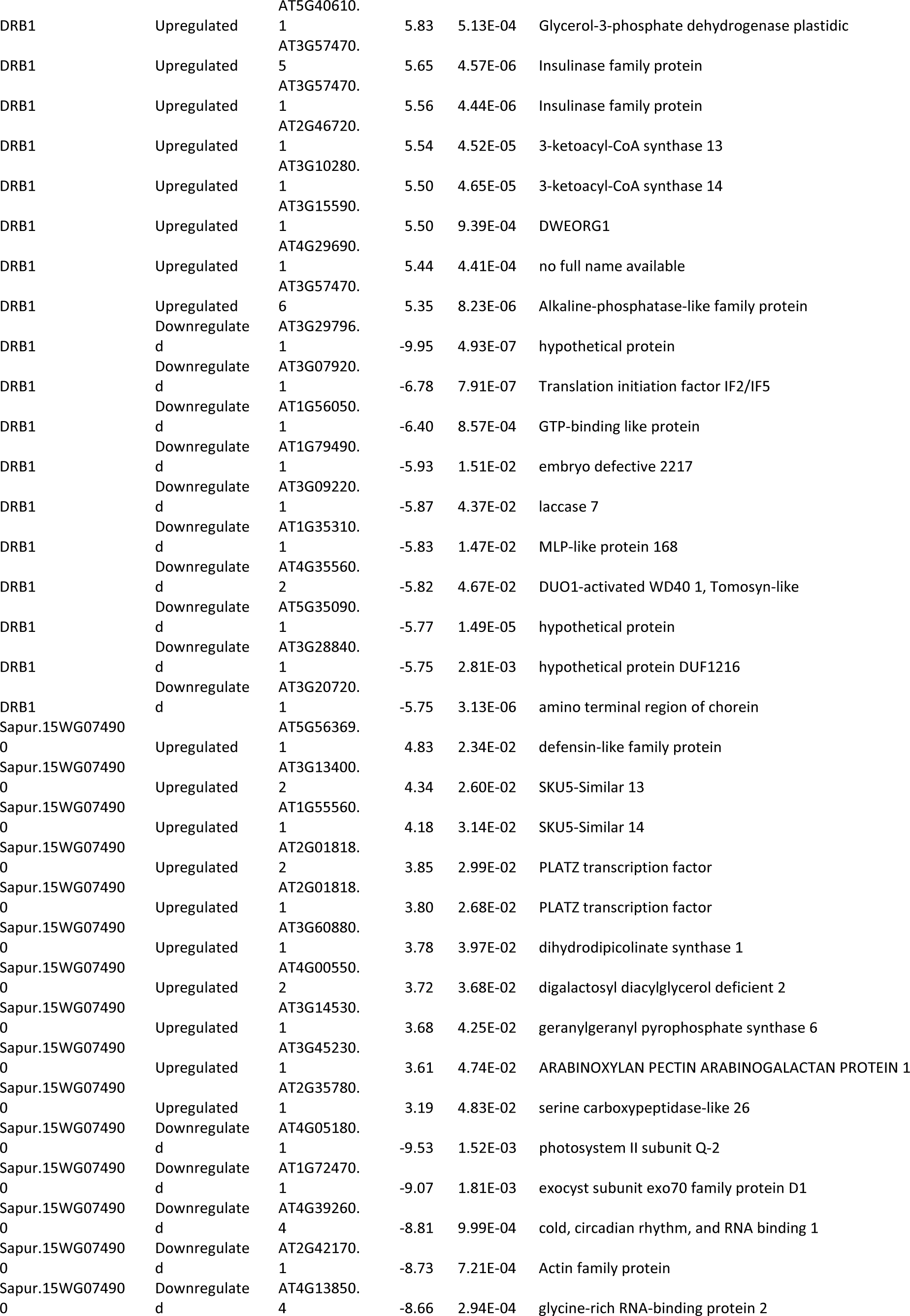
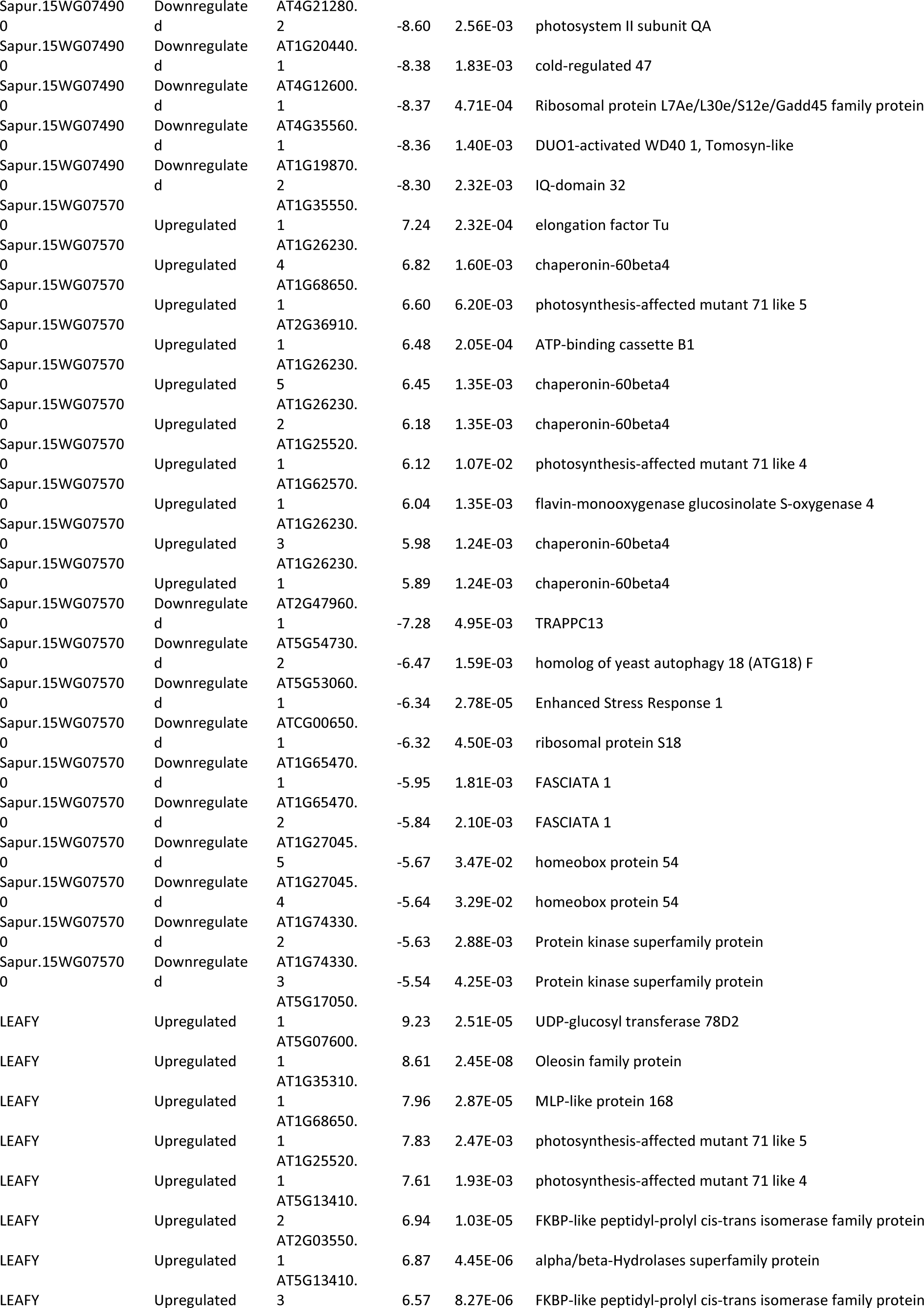
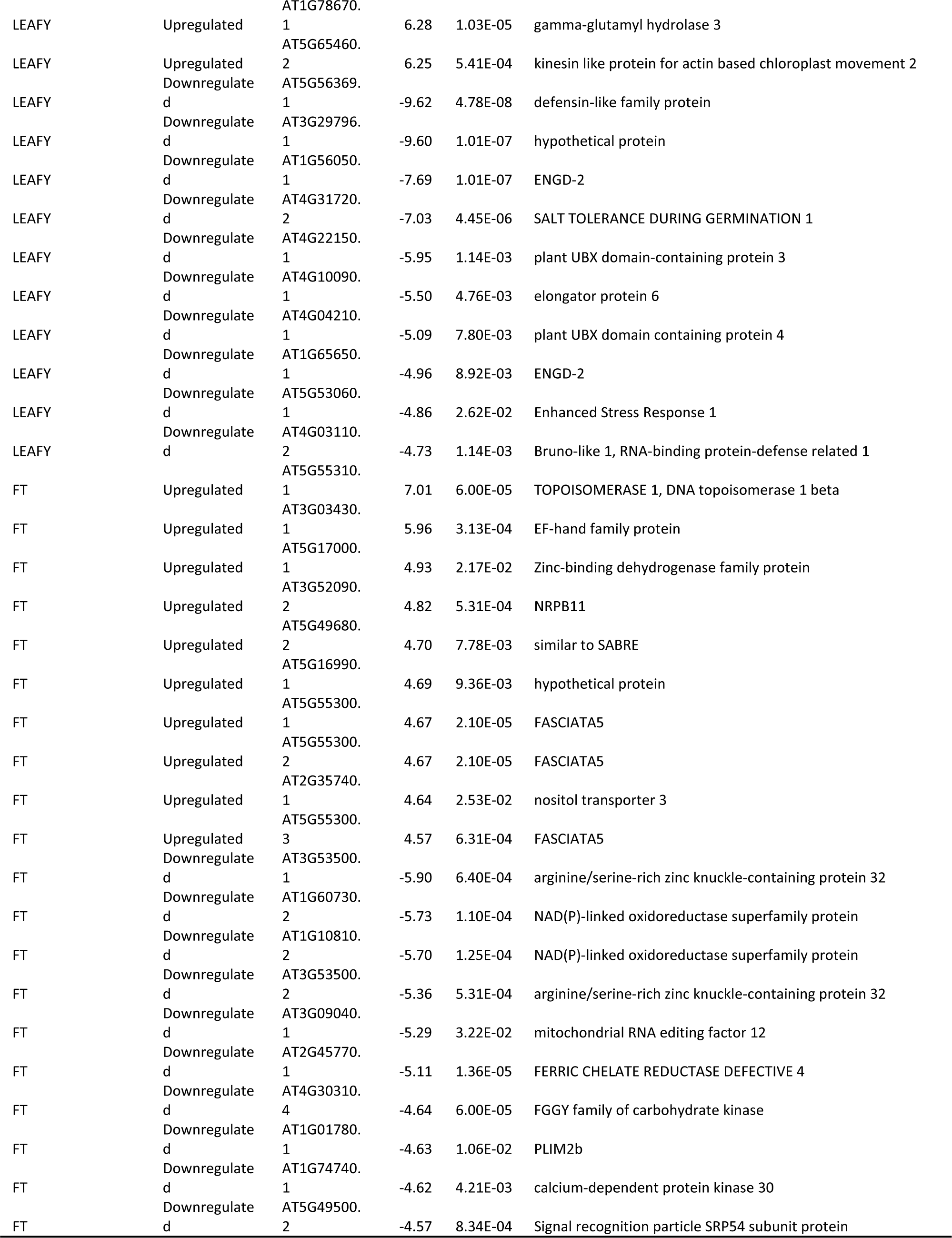
Top 10 greatest up- and down-regulated proteins for each Arabidopsis expression line.

**Table S2.**
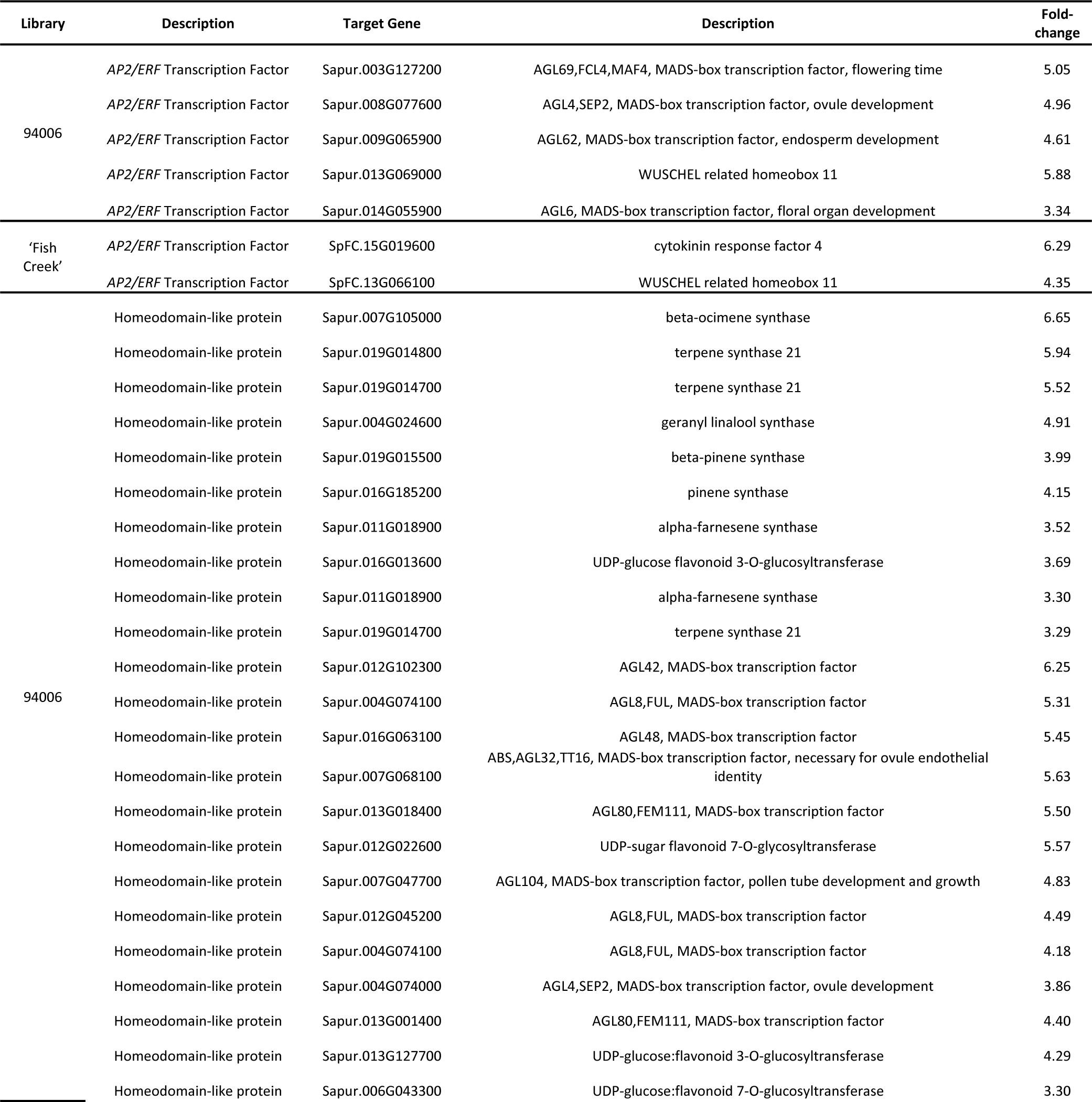
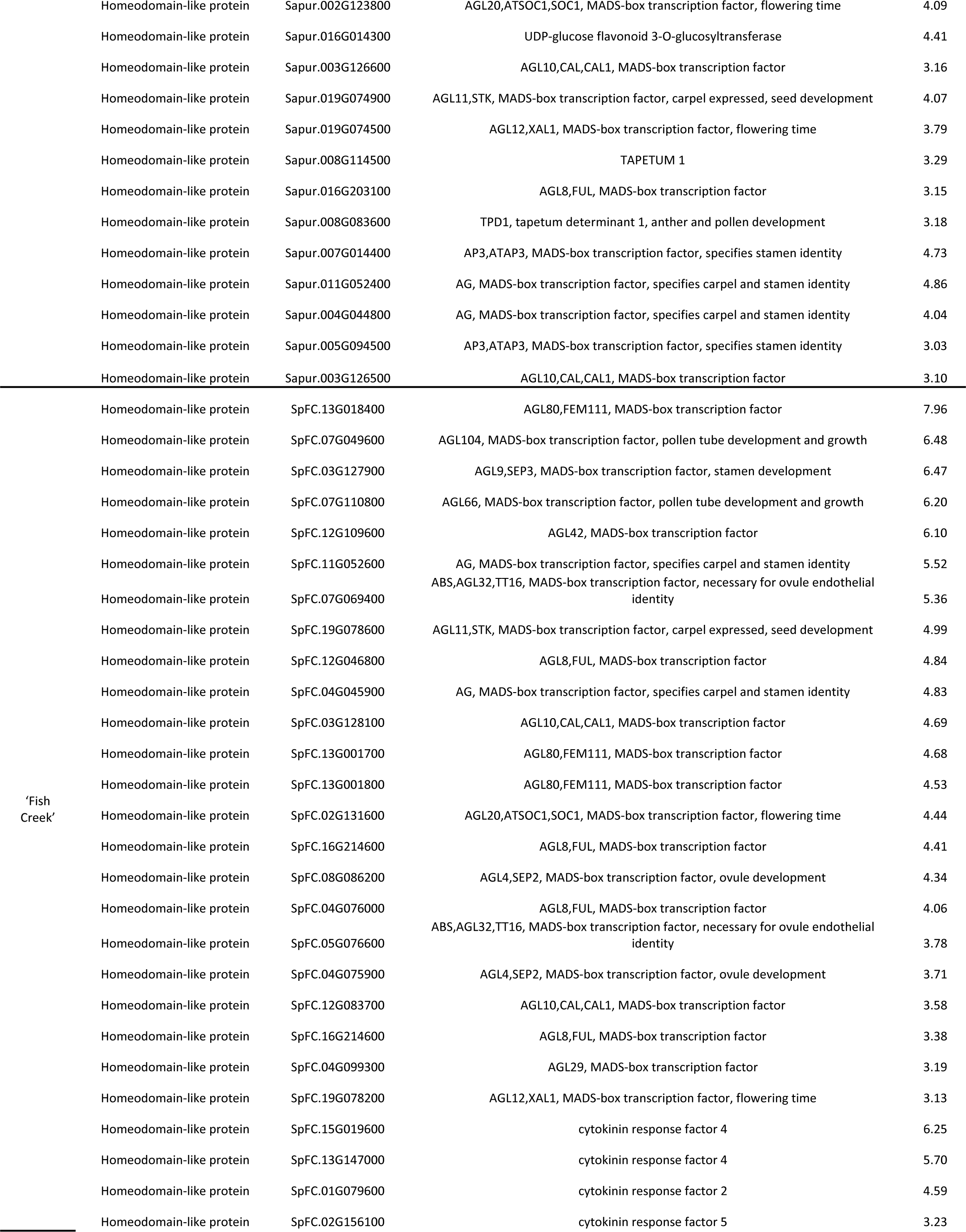
Sex dimorphism related genes involved in floral development and secondary metabolism adjacent to significant peaks in DAP-Seq analysis.

